# Protective role of the Atg8 homologue Gabarapl1 in regulating cardiomyocyte glycophagy in diabetic heart disease

**DOI:** 10.1101/2021.06.21.449174

**Authors:** Kimberley M. Mellor, Upasna Varma, Parisa Koutsifeli, Claire L. Curl, Johannes V. Janssens, Lorna J. Daniels, Gabriel B. Bernasochi, Antonia J.A. Raaijmakers, Victoria L. Benson, Eleia J. Chan, Marco Annandale, Xun Li, Yohanes Nursalim, Wendy T.K. Ip, David J. Taylor, Koen Raedschelders, Aleksandr Stotland, Aaron E. Robinson, Richard J. Mills, Regis R. Lamberts, Kim L. Powell, Terence J. O’Brien, Rajesh Katare, Chanchal Chandramouli, Rebecca H. Ritchie, Shiang Y. Lim, Robert G. Parton, Xinli Hu, James R. Bell, Enzo R. Porrello, James E. Hudson, Rui-Ping Xiao, Jennifer E. Van Eyk, Roberta A. Gottlieb, Lea M.D. Delbridge

## Abstract

Diabetic heart disease is highly prevalent and characterized by diastolic dysfunction. The mechanisms of diabetic heart disease are poorly understood and no targeted therapies are available. Here we show that the diabetic myocardium (type 1 and type 2) is characterized by marked glycogen elevation and ectopic cellular localization - a paradoxical metabolic pathology given suppressed cardiomyocyte glucose uptake in diabetes. We demonstrate involvement of a glycogen-selective autophagy pathway (‘glycophagy’) defect in mediating this pathology. Genetically manipulated deficiency of Gabarapl1, an Atg8 autophagy homologue, induces cardiac glycogen accumulation and diastolic dysfunction. Stbd1, the Gabarapl1 cognate autophagosome partner is identified as a unique component of the early glycoproteome response to hyperglycemia in cardiac, but not skeletal muscle. Cardiac-targeted *in vivo* Gabarapl1 gene delivery normalizes glycogen levels, diastolic function and cardiomyocyte mechanics. These findings reveal that cardiac glycophagy is a key metabolic homeostatic process perturbed in diabetes that can be remediated by Gabarapl1 intervention.

## Introduction

Diabetic heart disease incidence is increasing, associated with higher prevalence of type 1 (T1D) and type 2 (T2D) diabetes, and is a condition for which there is no specific therapeutic. Diabetic heart disease is recognized as a distinct cardiomyopathy^1^, characterized by early diastolic dysfunction^2^. Whilst cell signaling disruption associated with various aspects of diabetic cardiac pathology has been described^3–8^, the underlying mechanisms which drive the occurrence and fatal progression of diastolic dysfunction in diabetes remain poorly understood. Insulin deficiency (T1D) and insulin resistance (T2D) are both linked with reduced cardiac glucose uptake, although changes in insulin signaling pathway intermediates are variable depending on severity and duration of disease^9, 10^.

Altered myocardial substrate metabolism in diabetes has been reported with a shift from glycolytic fuel (glucose) to enhanced fatty acid utilization^11, 12^. Increased cardiomyocyte lipid storage and lipotoxicity are well described features of diabetic heart disease^13, 14^. Investigation of the metabolic and structural aspects of glucose storage in the form of glycogen fuel depots is surprisingly limited.

Glycogen reservoirs have been conventionally considered to provide a short term physiologic endogenous energy ‘buffer’ – a fuel reserve to maintain performance in settings of acute metabolic stress. However, some evidence suggests that glycogen may have a more dynamic and influential role in myocardial metabolic health^15–17^. Post-mortem clinical histologic evidence of cardiac glycogen accumulation in diabetic myocardium was first reported anecdotally 90 years ago^18^ – yet surprisingly this phenomenon has not been pursued. Impaired cardiomyocyte glucose uptake in diabetes would be expected to be associated with a corresponding decrease in glycogen levels, as is observed in non-cardiac tissues^19–22^. However findings gleaned from studies where cardiac glycogen has been incidentally evaluated are equivocal (as reviewed in Varma et al. 2018 ^8^). Despite the well-established literature identifying cardiac vulnerability in inherited glycogen storage diseases^23, 24^, cardiac glycogen storage dysregulation in diabetes has not been systematically examined.

Autophagy, a cellular macromolecular ‘bulk’ catabolic pathway, has been shown to be involved in progression of heart failure^12, 25–28^. Macro-protein and mitochondrial autophagy have been most well described, involving ‘cargo’ tagging by p62 (a ubiquitin-binding adaptor) which complexes with LC3B (an Atg8-family partner protein) to capture the cargo in an autophagosome destined for lysosomal fusion and subsequent macromolecule degradation. Findings regarding autophagic contribution to diabetic heart disease progression are discrepant^25, 27^.

Recently, a new understanding of selective autophagy pathways has emerged^29^. Sequence analysis has revealed that the glycogen-tagging protein, starch binding domain protein 1 (Stbd1), contains several binding motifs for Atg8 family members. In a non-cardiac cell line, the Atg8 homologue γ-aminobutyric acid receptor-associated protein-like1 (Gabarapl1, ambiguously historically named) was identified as a potential cognate binding partner for Stbd1^30^. Our preliminary cardiomyocyte *in vitro* studies have shown that Stbd1 and LC3B do not co-localize, indicating that Stbd1-Gabarapl1 processes may function in parallel with macro-autophagy p62-LC3B processes^31^. We postulate the operation of an autophagy pathway in the myocardium specific for glycogen (i.e. ‘glycophagy’).

Given the historical observation of glycogen deposition in human diabetic myocardium, the dramatic cardiac consequences of glycogen overload in glycogen storage diseases, and our preliminary findings indicative of a glycogen-selective autophagy route, the goal in this study was to delineate the role of glycophagy in the heart. We investigated the hypothesis that deranged glycophagy is an underlying metabolic defect in diabetic heart disease and is linked with cardiac functional deficit. Here we show that the diabetic myocardium is characterized by marked glycogen elevation and identify Stbd1 as a unique component of the early glycoproteome response to glycemic challenge in cardiac (but not skeletal) muscle. We demonstrate that Gabarapl1 plays an essential role in determining glycogen homeostasis and preserving diastolic function in the heart. Importantly we show that cardiac-specific *in vivo Gabarapl1* gene delivery normalizes glycogen levels and alleviates diastolic dysfunction in diabetes. Our work establishes glycophagy as a myocardial homeostatic process, which is deranged in diabetes and may represent a potential therapeutic target for cardiac functional remediation.

## Results

### Cardiac glycogen is linked with diastolic dysfunction in diabetes

In order to establish the status of cardiac glycogen levels in a range of diabetic settings, a comprehensive survey of glycogen content in myocardial tissues of diabetic patients and mature cardio-metabolic animal models was undertaken, including dietary, genetic, and pharmacologically induced diabetic states. This investigation revealed that in both T1D and T2D diabetic conditions, elevation of myocardial glycogen levels relative to non-diabetic controls was a consistent feature, with varying levels to a maximal 4 fold increase (Figure 1a). These findings represent a metabolic anomaly – in a context where cardiac glucose uptake is limited either by insulin deficiency or resistance, accumulation of glucose in glycogen polymer form seems paradoxical. We next examined the intracellular morphology associated with increased glycogen. A single branched glycogen molecule may comprise up to 120,000 glucose residues, and multiple molecules can be linked rendering these structures cytosolically identifiable using electron microscopy. In the myocardium, glycogen deposition was visualized as electron-dense particulates on transmission electron micrographs (optimized for glycogen opacity). In control hearts, glycogen particles were sparse. In contrast, marked glycogen infiltration was evident in diabetic myocardium, clustered near mitochondria and ectopically dispersed throughout the myofilament architecture (Figures 1b and S1a). To determine the cardiac specificity of the glycogen increase, we analyzed glycogen content in skeletal muscle tissue samples (quadriceps) collected in tandem with cardiac samples of several contrasting rodent diabetic models. Skeletal muscle glycogen levels in established diabetic models were not different to control, confirming a unique cardiac modulated glycogen phenotype (Figure 1c).

**Figure 1.**
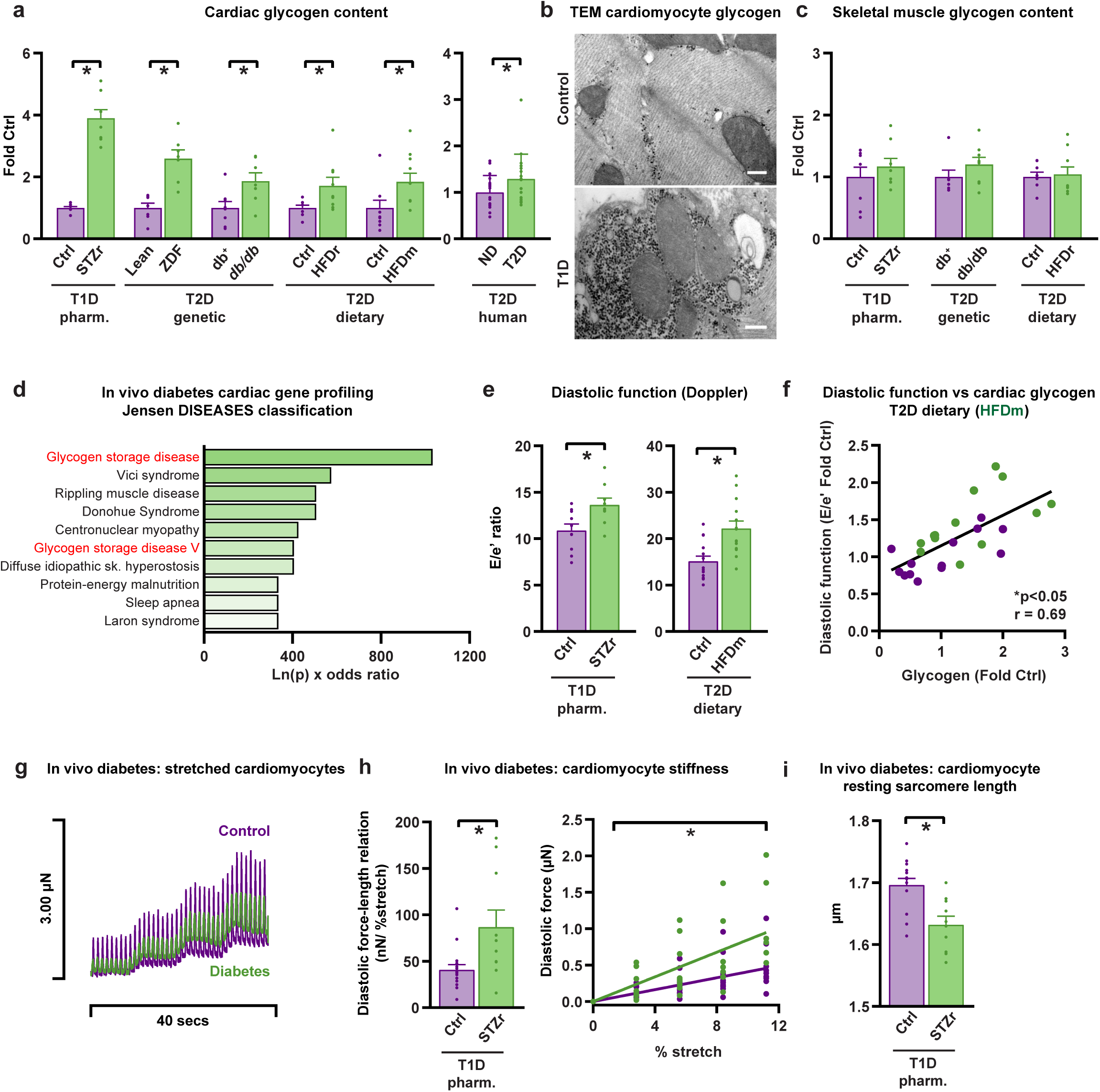
Cardiac glycogen is linked with diastolic dysfunction in diabetes. **a,** Cardiac glycogen content is increased in animal models of diabetes (T1D, type 1 diabetes; T2D, type 2 diabetes; mean ± SEM) and human patients with T2D (n=18-23 patients, mean ± SD). STZr, (streptozotocin-treated rat 8 weeks); ZDF (Zucker diabetic fatty rat age 20 week); db/db (leptin receptor mutant mouse age 10 week, db+ littermate control); HFDr (high fat diet-fed rat 14 weeks); HFDm (high fat diet-fed mouse 14 weeks), n≥7 animals. **b,** Diabetic cardiomyocyte transmission electron micrographs (TEM) of STZ rat heart show prominent electron-dense glycogen particulate deposits (black puncta) (scale bar 300nm). **c,** No change in skeletal muscle glycogen content in diabetic animal models (STZ rat; db/db mice, high fat diet rats (HFDr), n ≥ 7 animals). **d,** Jensen DISEASES classification analysis identified ‘glycogen storage disease’ as characteristic of differentially expressed genes detected by gene profiling diabetic rat hearts (STZ, n=6 rats, n=57 genes). **e,** Diastolic dysfunction in T1D rats (STZ) and T2D mice (HFD), demonstrated by increased echocardiographic index, E/e’ ratio (n≥10 animals). **f,** Myocardial glycogen content and diastolic dysfunction (E/e’ ratio) are correlated for control (purple) and diabetic (green, HFD) mice (r, Pearson correlation coefficient). **g,** Single intact cardiomyocyte force traces (exemplar) show higher diastolic force (an indicator of passive stiffness) in response to stretch in diabetic (STZ) vs control (Ctrl) rat. **h,** Cardiomyocyte end diastolic force-length relation (EDFLR) is increased in diabetic rats (STZ, n≥10 cells). Cardiomyocyte stiffness (diastolic force during sarcomere stretch) is increased in diabetic rats (green, STZ) relative to control (purple) rats (n≥10 cells). **i,** Cardiomyocyte resting sarcomere length is decreased in diabetic rats (STZ, n≥10 cells). Data presented as mean ± s.e.m. *p<0.05. See also Figures S1, S2, and Tables S1, S2.

To gain insight into the disease relevance of the glycogen phenotype, we performed gene profiling on ventricular mRNA samples extracted from diabetic rodents exhibiting the most marked elevation in glycogen (a T1D β-cell ablation model). The DISEASES gene association analysis tool^32^ was applied to a panel of 57 genes assembled to probe molecular signaling pathways involved in metabolic stress responses (Table S2). The differential gene expression profile was found to most closely align with a ‘glycogen storage disease’ state (Figure 1d).This is a large group of genetic conditions relating to a range of mutations involved in glycogen catalysis^23^.

We next investigated the activation state of the canonical cytosolic glycogen enzymatic regulatory pathway which controls the availability of free glucose by addition (synthase) or removal (phosphorylase) of single glucose units of glycogen branch structures. Glycogen synthase was found to be either inhibited (increased phosphorylation at Ser641) or unchanged, and glycogen phosphorylase was either activated (increased phosphorylation at Ser14) or unchanged, in a survey of diabetic rodent hearts (Figure S1b and S1c). Overall, these activation shifts would be consistent with suppressed glycogen synthesis and upregulation of cytosolic glycogen breakdown, certainly not instrumental as a causative mechanism of total myocardial glycogen elevation in diabetes. Based on these findings, alternative molecular pathways of glycogen management were implicated.

Given the known cardiac contractile deficits associated with diabetes, we next investigated whether a functional link with glycogen elevation could be identified in T1D and T2D settings. This investigation required matching echo data derived *in vivo* with analysis of tissues recovered post-mortem. Diastolic dysfunction *in vivo*, as typically clinically indexed by elevated ratio of the left ventricular blood/tissue Doppler signals (E/e’), was apparent in both T1D and T2D rodent models (Figure 1e). The elevated E/e’ levels recorded were commensurate with shifts considered clinically diagnostic of diastolic failure ^33, 34^. In T2D, no evidence of altered systolic function was detected, while small but significant reductions in ejection fraction and fractional shortening were observed in T1D animals where the diabetes phenotype was more progressed (Figure S2a-d). In the T2D model, where the data range was broader, a significant correlation between myocardial glycogen content and diastolic dysfunction was demonstrated (Figure 1f). An elevation of E/e’ as a measure of diastolic dysfunction defines the extent to which tissue is compliant during the early fast phase of ventricular blood inflow. A high value represents relative tissue stiffness. In order to evaluate the factors contributing to increased tissue stiffness, extracellular matrix fibrosis was quantified. Interstitial collagen (histologically assessed using picrosirius red staining) was increased in the T1D model but not in the T2D context (Figure S2e-f). Thus, in the more progressed diabetic conditions, increased fibrosis and systolic dysfunction were observed with diastolic dysfunction. In the T2D less functionally compromised states, diastolic dysfunction was present and correlated with glycogen, in the absence of systolic defect and without increase in collagen deposition. Whilst this finding would not preclude contribution of extracellular matrix alterations to tissue compliance, the possibility of non-matrix sources of tissue stiffness was suggested.

To determine the contribution of changes in intrinsic myocyte properties to *in vivo* diastolic dysfunction, cell mechanical stiffness was evaluated in single, intact enzymatically dissociated isolated cardiomyocytes derived from diabetic hearts. We developed methodology to apply an incremental stretch protocol to cardiomyocytes, normalized to achieve uniform sarcomere extension. Intact cardiomyocytes were attached via bio-adhesive material to glass microfibres for measurement of absolute force (diastolic and systolic) during programmed stretch protocols under loaded conditions with electrical stimulation to contract (Figure S2g-h). Diabetic cardiomyocytes exhibited increased mechanical stiffness, evidenced by elevated force response to a series of programmed length step changes, calculated to be comparative at sarcomere level. Exemplar records depict the steeper gradient of the diastolic force-length relationship in T1D cardiomyocytes (Figure 1g), with the mean normalized force-length relation increased more than 2-fold (Figure 1h). Basal increase in tonic stiffness was also apparent in diabetic cardiomyocytes, evidenced by reduced diastolic sarcomere length – indicative of relaxation defect at the myocyte level even in quiescence (Figure 1i). These data show that in diabetes, the diastolic dysfunction observed *in vivo* linked with myocardial glycogen accumulation, at least partially reflects altered cardiomyocyte nanomechanics.

Taken together, these findings establish that in both T1D and T2D, elevated cardiomyocyte glycogen is a generalized and paradoxical metabolic phenotype. We show that the extent of glycogen accumulation is correlated with diastolic dysfunction and is not explained by altered activation of canonical cytosolic glycogen handling enzymes. As gene expression profiling indicated, it could be considered that the cardiac diabetic metabolic condition presents as an acquired form of glycogen storage disease.

### Proteomic analyses identify glycophagy involvement in early glycogen dysregulation

Since cardiac and skeletal muscle glycogen responses in diabetes were markedly different, we compared the early hyperglycemia molecular signaling processes in these tissue types. To achieve an acute glycemic challenge (GC), rodents were treated with a single dose of streptozotocin to lesion pancreatic β cells to suppress insulin secretion, and thus effect a rapid increase in blood glucose. After 48 hours, blood glucose was elevated more than 50% in treated animals compared with controls (and relative to pre-treatment levels). Cardiac (left ventricle) and skeletal muscle (quadriceps) tissues were harvested. A significant elevation of glycogen level was apparent in cardiac, but not skeletal muscle samples (Figure 2a). Tissues were processed using serial centrifugation steps and maltodextrin elution to recover and enrich for the ‘glycogen proteome’ (i.e. glycogen-associated proteins). Proteomic analyses were performed using liquid chromatography and mass spectroscopy (LC-MS/MS). For each tissue type, proteins identified in control and glycemic challenge conditions were compared. In both cardiac and skeletal muscle extracts the majority of proteins were common to control and glycemic challenge conditions, with a smaller number of proteins uniquely detected in each condition. Overall, the cardiac glyco-proteome was more extensive than the skeletal proteome, comprising a larger number of total identified proteins (513 vs 366 proteins) – suggestive of increased complexity of the cardiac glycogen regulatory environment (Figure 2b).

**Figure 2.**
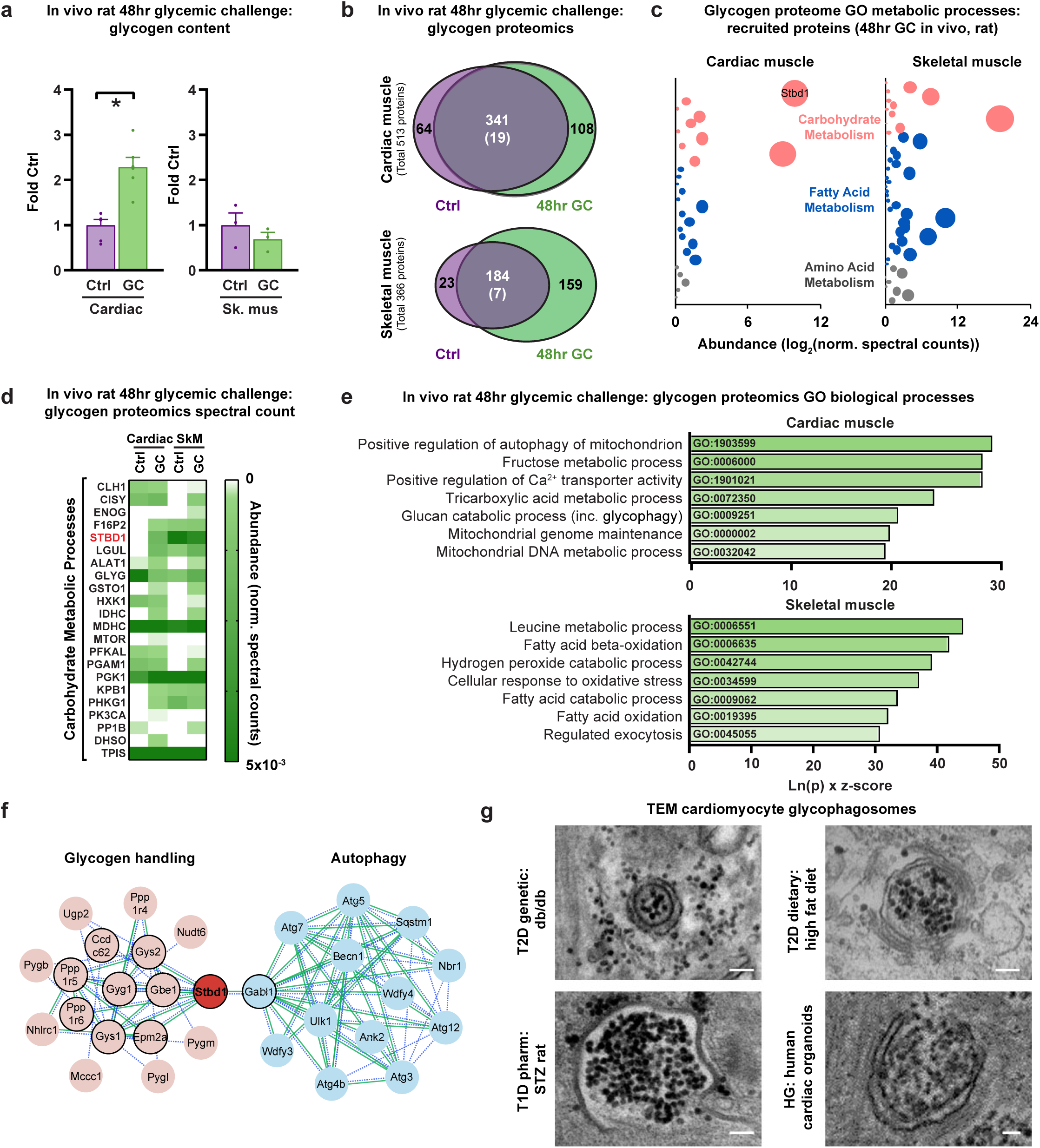
Proteomic analyses identify glycogen-autophagy (’glycophagy’) involvement in early glycogen dysregulation. **a,** Heart glycogen is increased in 48hr glycemically challenged (GC) rats and skeletal muscle glycogen is unchanged. The in vivo 48hr glycemic challenge was induced by 55mg/kg i.p. streptozotocin destruction of pancreatic b-cells (n=3-6 rats). **b,** Glycogen proteome in cardiac and skeletal (quadriceps) muscle after a 48hr glycemic challenge (GC) in vivo. Venn diagram values show the number of unique or differentially abundant proteins in control (Ctrl) vs 48hr GC samples using LC-MS/MS. Total number of significantly up- or down-regulated proteins shown in brackets. n=4 rats. **c,** The glycogen proteome response to a 48hr glycemic challenge is characterized by changes in proteins related to carbohydrate and amino acid metabolism in cardiac muscle, contrasting with skeletal muscle where changes in fatty acid metabolism related proteins are most evident. Data are derived from spectral counts of uniquely present proteins in 48hr GC rat tissue from Carbohydrate (GO:0005975), Fatty Acid (GO:0006629) or Amino Acid (GO:0006520) Metabolic Process GO categories, detected by LC-MS/MS (n=4 rats). Bubble size is indicative of protein abundance. **d,** The glycophagy tagging protein, Stbd1 (starch-binding domain-containing protein 1) is detected as a component of the cardiac glycogen proteome only in GC cardiac muscle (but not in control), whereas Stbd1 is present in both GC and control skeletal muscle glycoproteome. The heatmap is derived from LC-MS/MS normalized spectral counts of differentially abundant or uniquely detected proteins in the Carbohydrate Metabolic Processes GO category (GO:0005975), with normalized spectral abundance factor adjustment (n=4 rats). **e,** Glycophagy is identified as a key biological process in the cardiac glycogen proteome response to a 48hr glycemic challenge (GC) in vivo. The gene ontology (GO) categories with the highest combined score (Ln(p) x z-score) are presented, comprising glycoproteome proteins uniquely detected or differentially abundant between 48hr GC and control for heart and skeletal muscle (n=4 rats). **f,** Functional analysis network (STRING) of known and predicted protein interactions with Stbd1. Gabarapl1 is identified as a primary interactor with Stbd1 and a link between glycogen handling and autophagy. Primary interactions are depicted by black outline. Associations are derived from experimental evidence (continuous green line) and text mining evidence (dashed blue line). **g,** Transmission electron microscopy (TEM) images showing glycogen-enclosing autophagosomes detected in fixed myocardial tissues from rodent models of diabetes (type 2 diabetes (T2D), db/db mouse; type 1 diabetes (T1D), STZ treatment rat 8 week; T2D dietary, high fat diet rat), in human pluripotent stem cell-derived (hPSC) cardiac organoids cultured in high glucose (HG), scale bar 100nm. Data are presented as mean ± s.e.m. *p<0.05. See also Figures S3 and Table S3.

For each tissue, amongst the common proteins detected in both control and glycemic challenge conditions a relatively small number showed differential abundance (cardiac n=19, skeletal n=7). More notable, the response to glycemic challenge involved recruitment of unique components to the glyco-proteome in both tissues. To interrogate the metabolic characteristics of these tissue-specific responses to glycemic challenge we searched the group of tissue-specific recruited proteins using the Gene Ontology terms for Carbohydrate, Fatty Acid and Amino Acid Metabolic Processes (GO:0005975, GO:0006629, GO:0006520). The glycemic challenge profiles generated for each tissue type were markedly different (Figure 2c), with cardiac muscle proteins showing carbohydrate dominance, while fatty acid metabolism-related proteins were more prominent in skeletal muscle. Given the recognized role of lipotoxicity in diabetes-related cardiac disease, this finding was unexpected and indicates that myocardial carbohydrate metabolic derangement is an early response in glycemic dysregulation.

Remarkably, the most abundant cardiac metabolic protein unique to glycemic challenge hearts was the glycogen-tagging protein, Stbd1 (as annotated Figure 2c). As molecular analysis has shown that Stbd1 contains binding motifs for the autophagy Atg8 protein Gabarapl1, this protein could be considered functionally as an autophagy ‘adaptor’ for glycogen cargo. This is consistent with our finding that an Stbd1-glycogen association occurred as an initial response component specifically in cardiac muscle involved in the glycogen proteome remodeling in response to glycemic challenge. For the pooled group of cardiac and skeletal glycoproteome proteins uniquely recruited by glycemic challenge or differentially abundant in the ‘carbohydrate metabolic processes’ GO category, a map was constructed to compare control and glycemic challenge treatment values directly against matching skeletal muscle proteins (Figure 2d, Table S3). Amongst the 22 proteins involved, Stbd1 was distinctive – under control conditions it was robustly present in the skeletal glycoproteome, but not detectable in cardiac samples. Further, the responses to glycemic challenge were divergent as Stbd1 was recruited to the cardiac glycoproteome while levels were simultaneously diminished in the skeletal glycoproteome. Finally, a GO analysis was undertaken (Figure 2e) to glean higher order functional impacts of glycemic challenge for each tissue using the full sets of differentially abundant and uniquely detected proteins in both treatments (i.e. cardiac n = 64 + 19 + 108 = 191; skeletal n = 23 + 7 + 159 = 189). The cardiac response was characterized by changes in autophagy and glucan catabolic processes in concert with mitochondrial involvement. In contrast, the highest ranked processes in the skeletal muscle response involved fatty acid catabolism associated with oxidative stress. Thus, very different glycoproteomic metabolic responses were observed in different tissues of the same animals exposed to identical *in vivo* systemic acute glycemic challenge.

Collectively, these proteomic investigations targeting the earliest phase of systemic glycemic disturbance reveal that cardiac and skeletal glycogen management processes are fundamentally different. The construction of the glycoproteomes in each tissue reflects summation of disparate metabolic processes and impact on divergent biological processes. In response to an acute *in vivo* glycemic challenge, skeletal muscle maintains stable glycogen levels while cardiac glycogen paradoxically accumulates. This metabolic response pattern in the acute setting, associated with elevation of myocardial glycogen even during this short period, is consistent with the observed chronic state of glycogen accumulation in diabetic heart disease and suggests potential for disease causation. Stbd1 emerges as a crucial point of difference in determining the unique cardiac-patho-phenotype in the initial response to deranged systemic metabolism.

### Proteomic analyses link with ultrastructural evidence of glycogen-selective autophagy

Given these Stbd1 observations, we next performed a functional network analysis to probe the nature of the relationship between glycogen handling and linked effector proteins of importance in glycemic challenge pathology. The STRING database (Search Tool for Retrieval of Interacting Genes/Proteins) captures experimental evidence and performs text-mining to identify evidence of direct and indirect protein-protein associations in any organism setting. Using Stbd1 as the input protein, an association network was generated and depicted (Figure 2f) to illustrate primary (direct, black outline) and secondary protein interactions and designated links of specific evidence type. From the network generated, the Atg8 protein, Gabarapl1, emerged as the only autophagy protein with direct link to Stbd1 indicating that this pivotal Atg8 protein may provide the key nexus via Stbd1 as a driver of a glycogen-autophagy functional association, predictive of a specific cellular glycophagy apparatus.

These findings prompted us to seek ultrastructural evidence of myocardial glycophagy occurrence in diabetic contexts. We and others have previously noted autophagosomal punctate inclusions in non-disease settings^35, 36^. Given the disturbance of myocardial glycogen levels in diabetes we hypothesized that cardiac autophagosomal glycogen content could be increased. Using optimized methods of electron microscopy for glycogen visualization we were able to identify occurrence of glycogen-replete autophagosomes, which could be considered to reflect cargo-selective glycophagy activity. These structures were identified in rodent models of T1D and T2D, and in human pluripotent stem cell-derived cardiac organoids exposed to glycemic challenge (Figure 2g and S3).

Considered together, these proteomic, bioinformatic and ultrastructural findings provided evidence of the *in vivo* operation of a glycogen-selective autophagy route involving the adaptor-Atg8 scaffold partners (Stbd1 & Gabarapl1) distinct to the well recognized adaptor-Atg8 scaffold partners involved in general macroprotein autophagy (p62 and LC3B). Specifically in the myocardium (and not in skeletal muscle), we found Stbd1 was prominently recruited to the glycoproteome during an *in vivo* stress condition associated with cardiac glycogen accumulation, and that a unique Stbd1-Gabarapl1 association could be identified.

### Glycophagosome protein Gabarapl1 involvement in cardiac glycophagy flux disturbance in diabetes

As a next step we sought evidence for cardiac-specific glycophagic involvement in the pathologic glycogen response in diabetes. Based on our findings above and the general understanding of phagosomal mechanisms, the process of glycophagy can be understood to involve several stages – adaptor Stbd1 attachment to glycogen (tagging) and binding to Gabarapl1 at the forming autophagosome membrane, membrane enclosure of glycogen cargo, and subsequent fusion with a glucosidase-containing lysosome (acid α-glucosidase, Gaa) to degrade glycogen thereby releasing free glucose for local metabolic use. Thus, in parallel with the well characterized and regulated glycogen degradation by phosphorylase, glycophagy constitutes a ‘bulk’ mechanism of glucose release (Figure 3a). While the physiologic complementarity of these two routes of glucose supply could offer metabolic flexibility, we hypothesized a pathophysiologic role for defective glycophagy operation in diabetic glycogen dysregulation.

**Figure 3.**
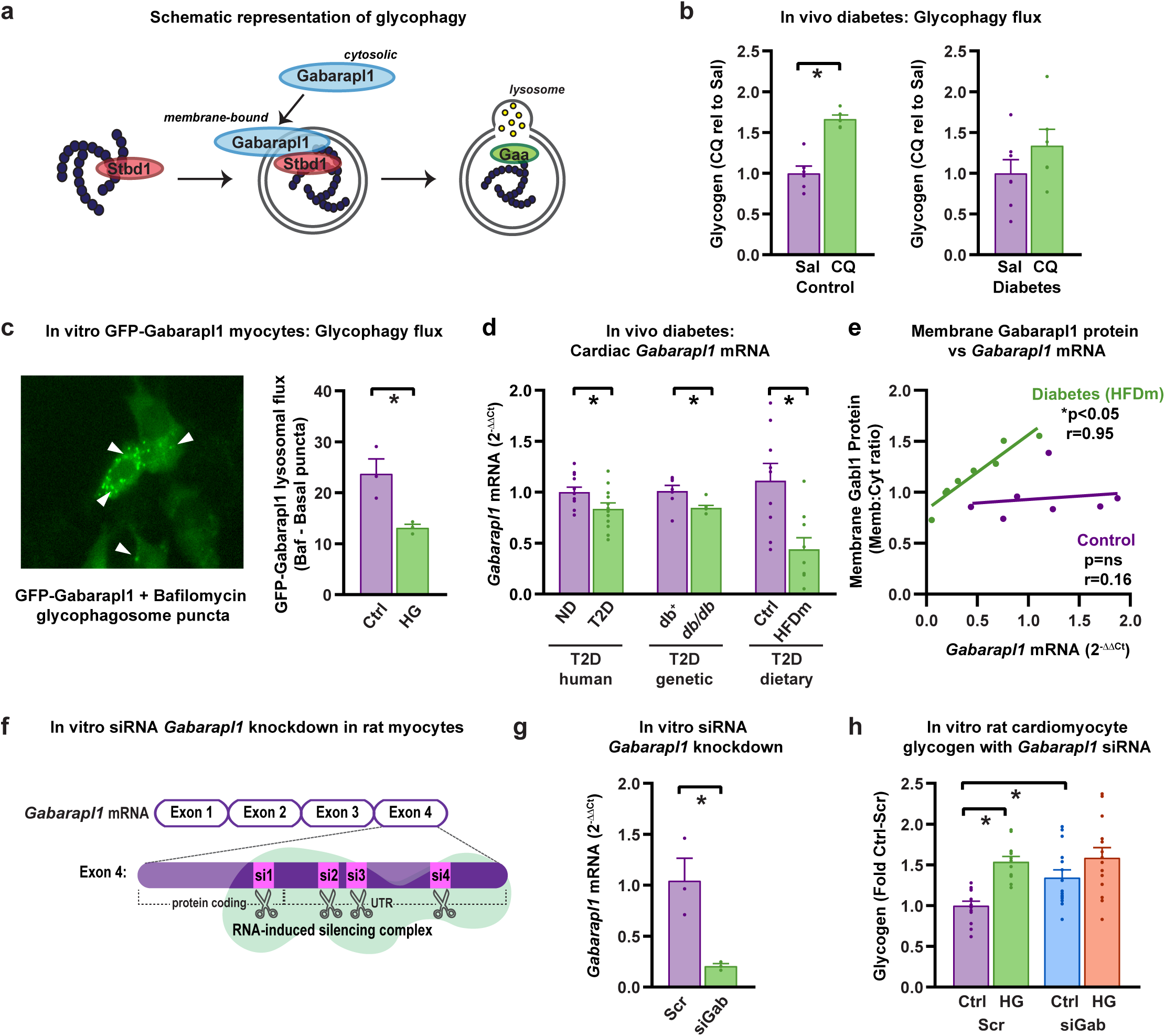
Glycophagosome protein (Gabarapl1) involvement in cardiac glycophagy flux disturbance in diabetes. **a,** Schematic illustrating three stages of glycophagy. Stbd1 tags glycogen and binds to the Atg8 protein, Gabarapl1 (GABA receptor-associated protein-like 1), at the forming autophagosome membrane. The mature glycophagosome fuses with a lysosome where Gaa (acid α-glucosidase) degrades glycogen to free glucose for metabolic recycling. **b,** In vivo glycophagy flux is impaired in diabetic hearts, shown as reduced glycogen accumulation with lysosomal inhibition (chloroquine (CQ), 50mg/kg i.p. 4 hours prior to tissue collection vs saline (Sal)) in diabetic rats (STZ, n=6-7 rats). **c,** Glycophagosome membrane-bound Gabarapl1 puncta visualized using GFP-Gabarapl1 in mouse atria-derived HL1 cardiomyocytes (white arrows). HL1 cells were transduced with GFP-Gabarapl1 and cultured in control (Ctrl) or high glucose (HG), imaged before (basal) and after 2 hours lysosomal inhibition with Bafilomycin (Baf). Gabarapl1 glycophagy flux is decreased in HG cultured HL1 cells, shown by reduced accumulation of GFP-Gabarapl1 puncta in response to lysosomal inhibition with Baf. Quantification of GFP epifluorescent micrographs presented as the number of puncta-containing cells as a percentage of total cells/60x image (9-13 images/well, n=3 independent wells). **d,** Cardiac mRNA expression of Gabarapl1 is decreased in human patients with diabetes mellitus (DM, n=12 patients), type 2 diabetic (T2D) hearts (db/db T2D mice with db+ littermate controls, age 10 week, n=7 animals; HFDm, high fat diet-fed mice 14 week, n=9 animals). **e,** Gabarapl1 mRNA is correlated with membrane-bound ‘active’ Gabarapl1 protein levels in diabetic mouse hearts (HFDm) but not control hearts, indicating that localization of Gabarapl1 protein to the glycophagosome is proportional to mRNA transcription signal in a diabetic setting (n≥7 mice; r, Pearson correlation coefficient). **f,** Schematic depicting small interfering RNA (siRNA) gene silencing experimental design using a pool of 4 siRNA sequences (si1-4) targeting Gabarapl1 for gene knockdown in neonatal rat ventricular myocytes (NRVM). **g,** Confirmation of siRNA-induced Gabarapl1 mRNA knockdown in NRVMs (n=3 independent wells). **h,** Cardiomyocyte glycogen content increased in NRVMs in response to high glucose (HG) or Gabarapl1 knockdown (siGab). No additive effect of HG and siGab observed, suggesting a common mechanism underlying glycogen accumulation in these settings (n=13-15 wells from 3 biologically independent cell culture experiments). Data are presented as mean ± s.e.m. *p<0.05. See also Figure S4.

To quantify cardiac glycophagy flux in diabetes *in vivo*, we measured the extent of glycogen accumulation in response to acute lysosomal inhibition in diabetic rodents (T1D, STZ-induced, 8 weeks duration). Autophagy is a dynamic process involving Atg8 protein movement in and out of phagosomes during the phagosome-lysosome cycle. Thus snapshot measures of total protein levels do not necessarily correlate with autophagy activity, whereas flux measurements provide a more informative indicator of phagosome throughput^37^. As previously established, acute chloroquine administration i.p. (with *in vivo* tolerability) arrests lysosome fusion and quantification of the accumulated Atg8 markers (e.g. LC3B) is an established method for monitoring macro-autophagy flux^37, 38^. By quantifying the glycophagy ‘cargo’, glycogen, following lysosomal blockade, we provide a method for monitoring glycogen-specific autophagic flux. As expected, chloroquine treatment increased cardiac glycogen in myocardium of control animals (Figure 3b). In diabetic animals where cardiac glycogen levels are elevated (Figure 1a), a blunted glycogen response to chloroquine was evident, indicating an already impaired cardiac glycophagy flux in diabetes (Figure 3b). Given that Stbd1 migration to glycogen appeared to be activated early in a glycemic challenge setting (Figures 2c and 2d), and phagosome glycogen localization in diabetic myocardium was apparent by transmission electron microscopy (Figure 2g and S3), we reasoned that impaired glycophagy flux could most likely involve defective phagosome processing, rather than phagosomal glycogen capture.

Next we used an *in vitro* approach to track cellular glycophagosome traffic in simulated diabetic conditions, employing a stable cardiomyocyte cell line expressing GFP-tagged Gabarapl1. The extent of Gabarapl1 puncta accumulation in response to bafilomycin-induced lysosomal inhibition (a specific inhibitor of vacuolar-type H^+^-ATPase in cultured cells^37, 39^) was blunted (Figure 3c), consistent with reduced glycophagy flux in diabetic milieu. In diabetic animal models *Gabarapl1* mRNA expression was significantly reduced, consistent with a reduction of *Gabarapl1* mRNA in diabetic human myocardium (Figure 3d). *Gabarapl1* mRNA levels were highly correlated with Gabarapl1 protein detected in the tissue homogenate membrane fraction (glycophagosome-localized) in diabetic, but not control samples (Figure 3e). These findings suggested that *Gabarapl1* mRNA availability was a limiting factor for glycophagy operation and supported the notion that targeted gene delivery of *Gabarapl1* could augment Gabarapl1 phagosomal localization and clearance.

To establish whether cardiomyocyte Gabarapl1 availability could be a determinant of cellular glycogen homeostasis, *in vitro* knockdown of *Gabarapl1* was performed in cultured neonatal rat ventricular myocytes using a pool of multiple siRNA constructs targeting the *Gabarapl1* gene (Figures 3f-g and S4a). *Gabarapl1* knockdown produced a marked increase in cardiomyocyte glycogen content under control incubation conditions (Figure 3h). Simulated diabetic incubation conditions (high glucose) induced an increase in cardiomyocyte glycogen content (Figure 3h). Combination of high glucose incubation with *Gabarapl1* siRNA did not produce an additive glycogen increase (Figure 3h), and suggested a common mechanism for glycogen accumulation induced by diabetic-like *in vitro* conditions and *Gabarapl1* gene knockdown. Collectively these *in vivo* and *in vitro* studies confirmed Gabarapl1 and glycophagy disturbance involvement in cardiac glycogen accumulation in diabetic conditions.

### Glycophagosome scaffold protein deficiency induces glycogen accumulation and diastolic dysfunction *in vivo*

Given the evidence that myocardial glycogen accumulation was linked with diastolic dysfunction in diabetes (Figure 1f), a cardiac-specific role for glycophagosome protein (Gabarapl1) deficiency in mediating glycogen disturbance and diastolic dysfunction was hypothesized. To determine the effect of Gabarapl1 deficiency on cardiac function and glycogen content, a Crispr/Cas9 global gene targeting approach was pursued to generate *Gabarapl1*-KO mice (Figure 4a and S4b-d). Heterozygote *Gabarapl1*-KO mice exhibited normal systemic and cardiac morphological characteristics with no change in body weight, glucose tolerance, blood glucose levels and heart weight (Figures 4b-d and S4e). Systolic function was preserved in heterozygote *Gabarapl1*-KO mice (ejection fraction, fractional shortening; Figures 4e-f), but diastolic dysfunction was evidenced by increased E/e’ (Figure 4g). The early report of cardiac autophagy by Ohsumi’s group^40^, identified the neonatal period of nutrition transition to be particularly reliant on autophagy activity. Thus, to assess the impact of *Gabarapl1* deficiency on myocardial glycogen levels, tissues of adult and neonate (age 2 days) mice were examined using enzymatic assay glycogen quantification. Further confirmation of glycogen deposition was sought in adult tissues using histological evaluation with periodic acid-Schiff staining. Evidence of glycogen accumulation was observed at both ages (Figures 4h and 4i). Taken together, these data provided evidence that *in vivo* Gabarapl1 deficiency is associated with impaired glycogen homeostasis linked with abnormal glycogen accumulation and cardiac diastolic dysfunction.

**Figure 4.**
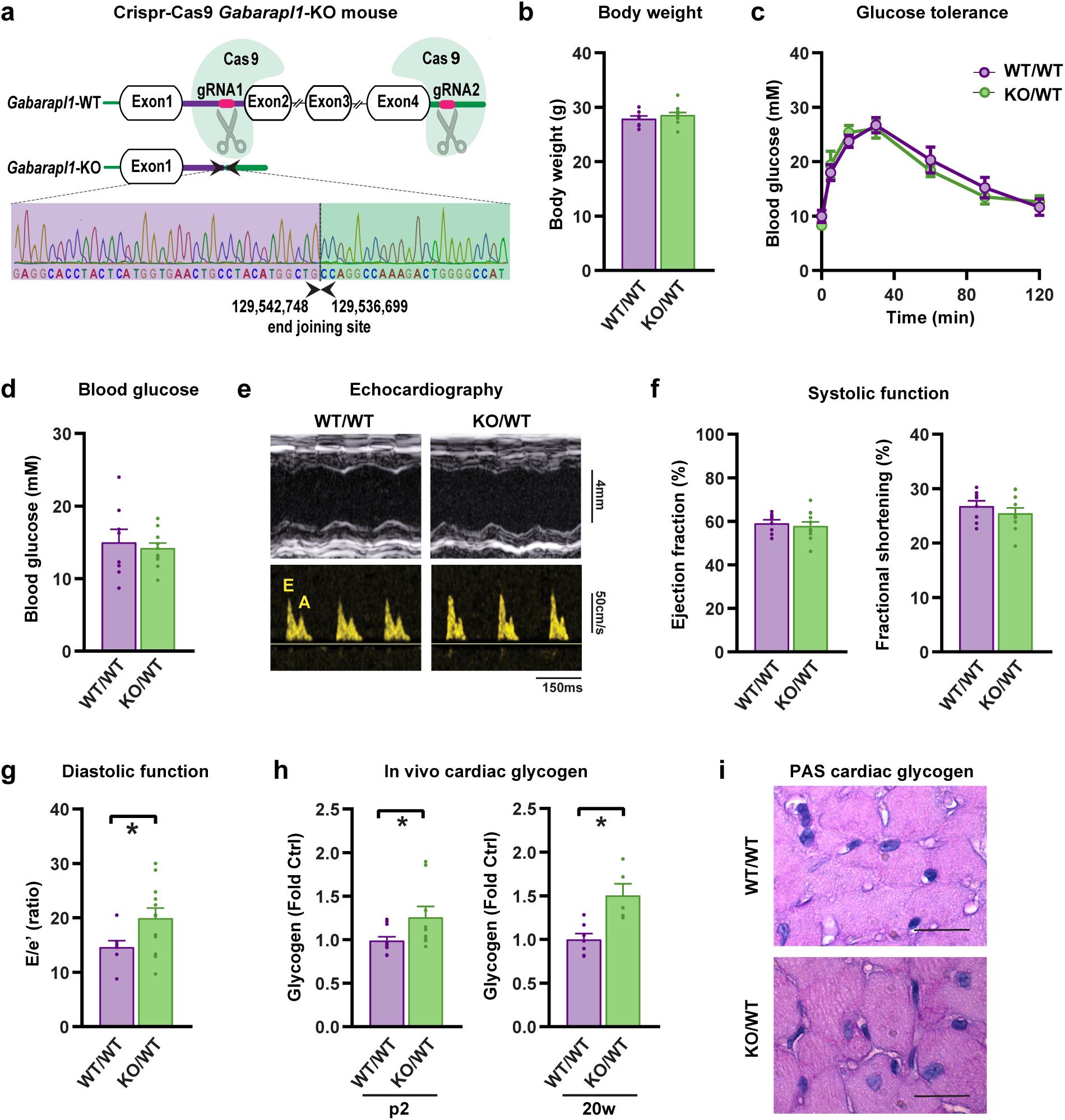
Glycophagosome scaffold protein deficiency induces glycogen accumulation and diastolic dysfunction in vivo. **a,** Schematic of Crispr/Cas9 genome editing design targeting Gabarapl1 for gene deletion in mice. Global Gabarapl1 knockdown did not affect **b,** body weight (n=8-12 mice), **c,** glucose tolerance (n=3-7 mice), or **d,** blood glucose levels (n=8-12). Despite no effect on systemic parameters, a cardiac phenotype was evident. **e,** M-mode echocardiography exemplar traces from left ventricular short axis view (upper panels) and pulse wave flow doppler (lower panels) in wildtype (WT/WT) and heterozygote Gabarapl1-KO mice (KO/WT). **f,** Systolic function was maintained in heterozygote Gabarapl1-KO mice (n=8-12). **g,** Heterozygote Gabarapl1-KO mice exhibit diastolic dysfunction, shown by increased E/e’ ratio (n=8-12 mice). **h,** Cardiac glycogen content is increased in neonate (p2, age 2 day) and adult (age 20 week) heterozygote Gabarapl1-KO mice (n=5-12). **i,** Periodic-acid Schiff (PAS) stained myocardial sections of 20 week old heterozygote Gabarapl1-KO mice show increased glycogen (pink-purple) staining (scale bar 50μm). Data presented as mean ± s.e.m. *p<0.05. See also Figure S4.

### Glycophagosome Atg8 gene therapy rescues cardiac glycogen accumulation and diastolic dysfunction in diabetes

To establish whether cardiac-specific augmentation of the expression of Gabarapl1 could mitigate progression of glycogen accumulation and diastolic dysfunction in diabetes, an AAV9-cTnT-*Gabarapl1* (AAV-Gab) viral overexpression approach (Figure 5a) was employed with rodent and human origin cells and tissues, and using an *in vivo* treatment for mice with established diabetic phenotype. In neonatal rat ventricular myocytes, high glucose-induced glycogen accumulation was prevented by transduction with AAV-Gab (Figure 5b and S5a). In human pluripotent stem cell-derived cardiac organoids transduced with AAV-Gab, the hyperglycemia-induced delay in force relaxation (diastolic dysfunction) was normalized with overexpression of Gabarapl1 (Figure 5c and S5b). In these PSC-cardiac organoids, systolic function was preserved under all conditions (Figure S5c).

**Figure 5.**
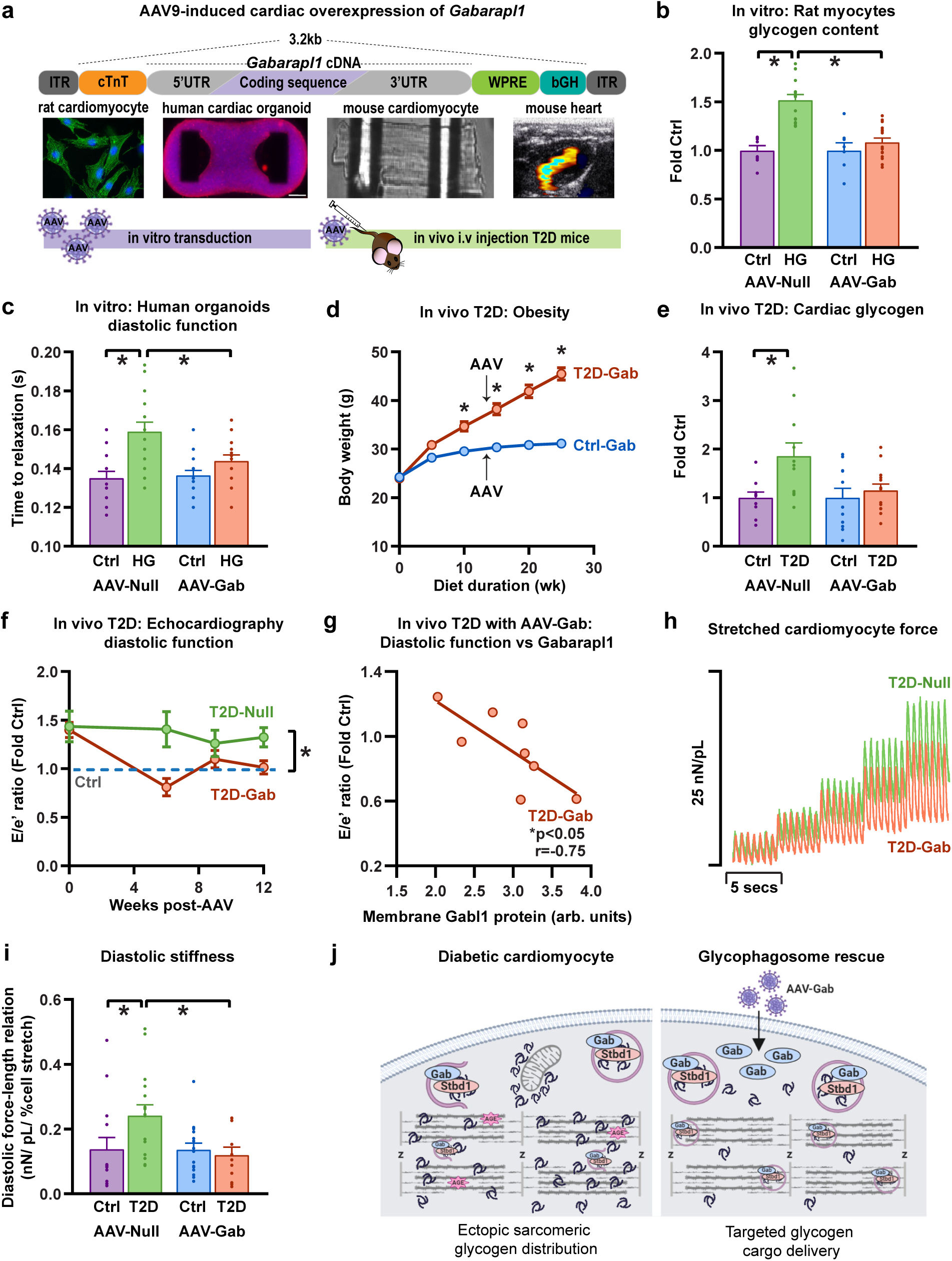
Glycophagosome Atg8 gene therapy rescues cardiac glycogen accumulation and diastolic dysfunction in diabetes. **a,** Schematic of cardiac-specific AAV9 gene targeting strategy involving rodent and human cell and tissue preparations and in vivo rodent interventions. **b,** High glucose (HG)-induced glycogen accumulation is rescued by AAV9-Gabarapl1 (AAV-Gab) transduction in cultured neonatal rat ventricular myocytes (n=9-11 independent wells, presented as fold change vs Ctrl-fed mice for each virus group). **c,** High glucose (HG)-induced diastolic dysfunction (delayed time to 50% relaxation of force) is rescued by AAV9-Gabarapl1 transduction in human pluripotent stem cell-derived cardiac organoids (n=13-19 organoids). **d,** In vivo cardiac-specific AAV9-cTnT-Gabarapl1 gene delivery does not affect the development of obesity in T2D mice (high fat diet, n=10 mice). **e,** T2D-induced cardiac glycogen accumulation is not evident in mice with cardiac Gabarapl1 overexpression (13 weeks post-AAV injection, high fat diet, presented as fold change vs Ctrl-fed mice for each virus group, n=10-12 mice). **f,** T2D- induced diastolic dysfunction (E/e’ ratio, echocardiography) is rescued by cardiac Gabarapl1 overexpression in AAV-transduced mice (high fat diet, presented as fold change vs Ctrl mice for each virus group, n=9-13 mice). **g,** Diastolic function (E/e’) is negatively correlated with Gabarapl1 protein expression in the membrane-enriched fraction of protein homogenates from T2D mice (high fat diet) injected with AAV-Gab virus (12-13 weeks post-AAV injection, r, Pearson correlation coefficient). **h,** Single intact cardiomyocyte force traces (exemplar) show rescued diastolic force (an indicator of passive stiffness) with cardiac Gabarapl1 overexpression in diabetic mice (AAV9-cTnT-Gabarapl1; T2D, high fat diet; 24 weeks post-AAV injection). **i,** T2D-induced cardiomyocyte diastolic stiffness is rescued by cardiac Gabarapl1 overexpression in diabetic mice (end-diastolic force-length relation slope; isolated stretched cardiomyocytes from mice at 24 weeks post-AAV injection, high fat diet, n=11-16 cells). **j,** Proposed model of glycophagy rescue targeting diastolic dysfunction in diabetes. Impaired glycophagy in diabetic cardiomyocytes is associated with glycogen accumulation. Ectopic intramyofibrillar glycogen may elicit localized glucose toxicity and advanced glycation-end product (AGE) formation at myofilament sites. Cardiac gene therapy with Gabarapl1 promotes glycophagic degradation of glycogen and localized release of glycolytic fuel at sarcoplasmic reticulum terminal cisternae (z-disks). Schematic prepared using BioRender.com. Data are presented as mean ± s.e.m, *p<0.05. See also Figure S5 and Table S4.

To implement an *in vivo* Gabarapl1 ‘rescue’ intervention, AAV-Gab virus titres were injected by tail vein into T2D (high fat diet) and control mice after 14 weeks of dietary treatment (Figure S5d-e). Mice were monitored for an additional treatment period of 12 weeks. As expected, the cardiac-specific AAV-Gab virus had no effect on systemic measures of body weight and glucose tolerance (Figure 5d and S5f). T2D-induced cardiac glycogen accumulation was not evident in T2D mice treated with AAV-Gab (Figure 5e). Activation states of glycogen synthase and phosphorylase were unchanged with T2D and AAV-Gab (Figure S5g-h), indicating that the suppression of glycogen accumulation in T2D with AAV-Gab treatment was linked with upregulation of the glycophagy pathway. The macro-autophagy marker, LC3B (ratio of lipidated to un-lipidated form) was also unchanged (Figure S5i). Diastolic dysfunction in diabetic mice (evidenced by increased E/e’ in T2D-Null mice) was rescued in T2D mice injected with AAV-Gab by 6 weeks post-AAV9 injection, a differential that remained evident at 12 weeks (Figure 5f and S5j). E/e’ was negatively correlated with Gabarapl1 protein expression in the membrane-enriched fraction of cardiac protein homogenate (Figure 5g), suggesting that diastolic function improvement in AAV-Gab-treated T2D mice was linked with augmented incorporation of Gabarapl1 protein into the phagosome membrane. Systolic function was unaffected by diabetes and AAV9 treatment (Figure S5j and Table S4). Thus, cardiac-specific Gabarapl1 expression upregulation reversed myocardial glycogen accumulation and diastolic dysfunction in a model of diabetes progression.

To advance the gene therapy investigation to the cellular level, cardiomyocytes were isolated from T2D mice at 24 weeks after the AAV-Gab injection. Single cardiomyocyte nano-mechanical properties were evaluated as described earlier (Figure S2h). Consistent with rescued diastolic dysfunction evident *in vivo*, the slope of the end-diastolic force-length relation (an index of myocyte stiffness) was significantly increased with diabetes and restored to control levels with overexpression of Gabarapl1 (Figure 5i). Collectively, these data support the contention that an intervention promoting availability of the glycophagosomal Atg8 membrane scaffolding protein (via AAV-*Gabarapl1* gene therapy) protects against cardiomyocyte diastolic dysfunction and glycogen accumulation in diabetic heart disease (Figure 5j).

## Discussion

In this study we demonstrate that in the myocardium, glycogen metabolism via a glycogen selective form of autophagy (glycophagy) is operational, and that disrupted glycophagy plays a pathophysiologic role in diabetic heart dysfunction. Findings show that elevated myocardial glycogen levels are a specific feature of diabetic heart disease (T1D & T2D), involving cardiomyocyte and cardiac diastolic abnormality. Glycogen levels in skeletal muscle are not similarly perturbed in diabetes. A deficiency of available Gabarapl1, a glycophagosome Atg8 and binding partner of the glycogen tagging protein Stbd1, is linked with cardiac glycogen accumulation. Genetically induced Gabarapl1 deficiency *in vivo* recapitulates diabetic heart disease in the absence of systemic diabetic disturbance. In established diabetes, cardiac-specific *Gabarapl1* gene delivery normalizes glycogen content, diastolic dysfunction and cardiomyocyte mechanics. These findings provide compelling evidence that cardiac-specific regulation of glycophagy constitutes a major cardiomyocyte metabolic axis and exerts profound functional influence.

### Myocardial glycogen metabolism and localization

Various aspects of cardiomyocyte cytosolic glycogen enzymatic regulation have been described, primarily in relation to ischemic cardio-protection. The cardiomyocyte glycogen store is in constant turnover, facilitated by the combined actions of glycogen synthase, phosphorylase, branching and debranching enzymes and involving numerous regulatory signaling intermediates including AMPK, PI3K and GSK3β^23, 41^. Glycogen metabolism significantly contributes to myocardial ATP production^20, 42, 43^. Endogenously derived glucose (i.e. from glycogen) is preferred over exogenously supplied glucose as an oxidation substrate ^20^. Importantly, metabolism of glycogen-derived glucose is linked with optimal ‘metabolic coupling’ of glycolysis to glucose oxidation, which is understood to minimize cytosolic proton loading, protecting against acidosis and maintaining myofilament sensitivity to activator Ca^2+^ ^20^. The cardioprotective importance of mitochondrial pyruvate carriers in connecting glycolysis and glucose oxidation has been recently highlighted ^44–46^. Observations made many decades ago^47, 48^, showed that glycolysis-derived ATP is particularly important in supporting cardiomyocyte Ca^2+^ homeostatic processes integral to electromechanical coupling. Physiologically, glycogen metabolizing enzymes appear to co-localize with the sarcoplasmic reticulum (SR) within the sarcomere, optimally positioned to deliver glucose for glycolysis to fuel Ca^2+^ re-uptake into the SR during cardiomyocyte relaxation by the SR-Ca^2+^ ATPase^49, 50^. The metabolic machinery enabling this process of substrate targeting has not previously been identified.

Here, with a new understanding of glycogen phagosomal handling, we bring these metabolic perspectives together to postulate a novel model of glycogen trafficking in the cardiomyocyte (Figure 5j). It has been suggested that physiologically, exogenous glucose entering the cytoplasm is substantially directed into glycogen synthesis (perhaps to protect cell constituents against glycosylation)^48^. We postulate that when matured, multi-branched glycogen molecules are Stbd1-tagged and via Gabarapl1 association, are captured as glycophagosome cargo. This cargo could be tracked to both peri-mitochondrial and para-SR cisternae sites. At these delivery sites, disgorged ‘free’ glucose would then be strategically positioned for tightly coupled mitochondrial oxidation and/or for glycolysis support of the SR-Ca^2+^ ATPase and other ATP-requiring ion pumps involved in electromechanical coupling. The phagosome released Stbd1 and Gabarapl1 components would then be recycled for new cargo collections in this physiological glycophagy flux functional scenario.

On the basis of our findings, to further extrapolate, we suggest that glycophagy disturbance in diabetes disrupts targeted tracking and/or docking of glycophagosomes at delivery sites. It could be speculated that Stbd1-tagged glycogen incorporation into the glycophagosome is impaired due to relative Gabarapl1 deficiency. It is possible that this Atg8 protein has a dual role in forming phagosomes for targeted translocation of glycolytic substrate, and in docking phagosomes at glycolysis enzyme hubs or ‘anchor’ sites. When Gabarapl1 levels are limiting, some glycophagosomes may be simply immobilized (with Gabarapl1/Stbd1 entrapment) (Figure S4), and some may disintegrate mechanically. This would be consistent with glycogen ectopically dispersed through the myofilament architecture (Figures 1b and S1). Similarly, the observed excess of glycogen clustered near mitochondria (Figure 1b) indicates ultrastructural evidence of metabolic uncoupling, whereby peri-mitochondrial pyruvate supply for mitochondrial uptake and oxidation is impeded (and thus fatty acid oxidation is correspondingly increased). Little is known about management of phagolysosomal content efflux mechanisms, and this may be a new field of investigation particularly relevant to diabetic heart disease.

The phosphorylation activation states for glycogen phosphorylase (upregulated) and synthase (downregulated) measured in diabetic myocardium are consistent with increased local reliance on these enzymes for cytosolic glycogen degradation (Figure S2a and S2b) in circumstances where localized phagosomal glycogen breakdown is impaired. An additional issue with increased non-localized release of free glucose via cytosolic glycogen degradation could be production of a pro-glycation cellular environment. Elsewhere we have reported occurrence of advanced glycation end-product formation on myofilament proteins in the diabetic myocardium, consistent with increased exposure of myofilaments to free glucose glycating influence^51^. Speculative, yet plausible, this glycophagy narrative offers a first step forward in understanding cardiomyocyte glycogen trafficking mechanisms. We believe that our findings integrate a number of previously unconnected concepts relating to cardiac metabolism and function, and that the scene is set for new avenues of major investigation.

### The distinctive cardiac glycogen proteome

Our evaluation of the glycogen binding proteins aggregating in the early response to glycemic challenge reveals that very different processes are operative in cardiac and skeletal muscle. This characterization of the cardiac glycogen proteome provides a snapshot view of what has been previously hypothesized as a glycogen ‘anchoring’ complex^48^. The cardiac glycogen proteome is extensive. The 449 co-locating proteins in the early response to glycemic challenge indicates the complexity of metabolic regulatory hubs involved (compared with 343 proteins in skeletal muscle). Indeed the scale of both muscle proteomes is striking compared to the two other glycogen proteomes previously reported for hepatic and adipose tissues^52, 53^. Our earlier structural study of glycogen particulates indicates that ultra-purified glycogen particle dimensions are larger in heart than other tissues^54, 55^ suggesting that both the glycogen branch structure and the associated protein population add to glycogen particle bulk *in situ*.

Functionally most distinctive is the cardiac-specific behavior of Stbd1, which was not detected in the glycogen proteome under basal conditions and was the most robustly recruited protein in response to glycemic challenge (while the reciprocal shifts occurred in skeletal muscle). The pivotal link between Stbd1 and the Atg8 protein Gabarapl1 demonstrated by functional network analysis provided the impetus to interrogate the Gabarapl1 role in modifying glycogen handling in the chronic disease state.

### Diabetic heart disease metabolism – an acquired glycogen storage disease

Myocardial glycogen responses to systemic metabolic manipulations are anomalous. In an experimental fasting setting, myocardial glycogen increases, while skeletal glycogen decreases)^54, 56^. An acute bout of exercise elicits similarly disparate tissue responses^57, 58^. These cardiac glycogen shifts have been interpreted as metabolic ‘survival’ defense responses – a pre-emptive sequestration of potential ‘emergency’ fuel for a constantly active, high energy demand tissue. In contrast, it is also proposed that in failure, cardiac insulin resistance represents a metabolic defense to protect against fuel excess^48, 59^. While seeming contradictory, both these responses represent strategies to minimize overall cytosolic ‘free’ glucose levels, thus protecting against damaging glycation/glycosylation events.

The prominence of cardiac phenotype in various genetic glycogen storage diseases also reflects a specific tissue susceptibility. In some of these disease settings, interventions which unload the glycogen accumulation have been shown to produce functional gains. Clinically, Gaa gene therapy remediated cardiac dysfunction ^60^. Experimentally, glycogen storage disease induced by mutation of the AMPKγ2 subunit was partially rescued by inhibition of glycogen synthase^61^. Given the cardiac-specific and glycogen-selective Stbd1-Gabarapl1 disease axis we identified, and the Jensens ‘Glycogen Storage Disease’ classification assigned, we targeted Gabarapl1 for loss and gain of function investigations. Findings from our *in vitro* proof-of-concept experiments involving in cardiac cell lines, primary cardiomyocytes and human cardiac organoids, supported the rationale for *in vivo* longitudinal interventions.

Global deletion of Gabarapl1 (heterozygous) had no detectable effect on systemic parameters of glucose homeostasis or on body weight. Yet the cardiac phenotype was dramatic, with induction of marked diastolic dysfunction linked with glycogen accumulation. Conversely, results obtained from experiments using primary cardiomyocytes, human cardiac organoids and diabetic mice consistently showed that cardiac gene delivery of *Gabarapl1* is protective in regulating cardiomyocyte glycogen and rescuing diastolic dysfunction. We provide the first evidence that deletion of an Atg8-family protein induces diastolic dysfunction. Other Atg8 members could be involved in cardiac glycophagy, including Gabarap which has higher affinity for Stbd1 than LC3B^62^, and more exploration is required. Systemic/global autophagy interventions in diabetic rodent models have generated discrepant cardiac findings (reviewed in Delbridge *et al* 2017 ^25^), and surprisingly investigations of cardiac-specific manipulation of autophagy to ‘rescue’ established diabetic heart disease have been limited. Targeting Beclin1 (an upstream autophagy initiator) using a systemic TAT-peptide approach has provided some evidence of diabetic diastolic dysfunction remediation^63^. Given the generalized impacts of non-specific autophagy induction, it could be anticipated that more targeted specificity for autophagy subtypes such as glycophagy may deliver the most promising translational outcomes. An important caveat is that while our investigation does not definitively identify Gabarapl1 as the primary instigator of glycophagy dysfunction in diabetes, our findings clearly implicate Gabarapl1 mediation and show that a Gabarapl1-based intervention has efficacy.

### Glycogen, cardiomyocyte stiffness & diastolic dysfunction

At present there is limited understanding of how loss of myocardial compliance in cardiac disease may in part reflect altered cardiomyocyte stiffness. Our findings that glycogen accumulation is linked with single myocyte mechanical stiffness and correlated with diastolic dysfunction provide new insight. Ultrastructurally, ectopic localization of glycogen may cause mechanical/ physical disturbance to sarcomeric force transmission. Further, disrupted intracellular glucose flux exposes the contractile machinery to glycation, potentially affecting sarcomeric compliance through crosslink/adduct formation and altering relaxation by molecular modification of protein domains crucial in electromechanical transduction. The gradual accumulation of these deficits would be consistent with the long subclinical phase of diastolic dysfunction in diabetic heart disease.

In conclusion, our study has identified that the Atg8 homologue Gabarapl1 interacting with a cognate binding partner Stbd1 to mediate glycogen autophagy, comprises a key cardiac metabolic axis. The demonstration of functional glycophagy as a physiologic metabolic route, and dysfunctional glycophagy as a diabetic cardiac metabolic pathology offers potential for development of interventional strategies. Almost three decades since Ohsumi’s seminal autophagy discovery work^64^, these findings may contribute to achieving translational outcomes to remediate metabolism and function in diabetic heart disease.

## Acknowledgements

We acknowledge J Liu, and E.M. Stevens for technical assistance, the Biomedical Imaging Research Unit University of Auckland for electron microscopy technical support, the Melbourne Histology Platform and Australian Phenomics Network for assistance with histological processing, the Bio21 Institute of the University of Melbourne for mass spectrometry support, G. Nolan for providing Phoenix-GP cells for HL1 transduction, D. Stapleton for providing the Stbd1 antibody, the Melbourne Advanced Genome Editing Centre for generating Crispr-Cas9 mice, I. van Hout for curating the HeartOtago tissue bank database, and P. Davis and D. Parry for obtaining tissues from patients undergoing surgery at Dunedin Hospital.

This work was supported by grants from the National Health and Medical Research Council of Australia (NHMRCA;1027865, 628643, 1082215, 1037320, 1067869), the Diabetes Australia Research Trust, Stem Cells Australia, the National Heart Foundation of Australia (NHFA), the New Zealand Marsden Fund (14-UOA-160), the Health Research Council of New Zealand (19/190) and the University of Auckland Faculty Research Development Fund. Fellowship support is acknowledged from NHMRCA (JEH, RGP, ERP), and Heart Foundation of Australia (ERP, JEH).

## Author contributions

L.M.D.D and K.M.M conceived the project and designed the experiments. K.M.M., U.V., P.K., C.L.C., J.V.J, L.J.D, W.T.K.I, A.J.A.R, G.B.B, V.L.B, E.J.C., M.A., X.L., Y.N., D.J.T., K.R, A.S., R.J.M., R.G.P., and X.H. planned and performed experiments, and analyzed data. K.M.M., U.V., P.K., C.L.C., J.V.J, W.T.K.I, A.J.A.R, G.B.B, V.L.B, D.J.T., K.R, A.E.R, A.S., R.R.L, R.K, K.L.P., T.J.O., C.C., R.H.R., S.Y.L, X.H., E.R.P., J.E.H., R.P.X., J.E.V.E., R.A.G., L.M.D.D provided materials and/or methodologic input. K.M.M., U.V., P.K., C.L.C., J.V.J, L.J.D, G.B.B, V.L.B, X.H., J.R.B., E.R.P., J.E.H., R.P.X., J.E.V.E., R.A.G., L.M.D.D provided interpretative review of study findings. K.M.M., U.V., P.K., J.V.J, L.J.D, E.R.P., J.E.H., R.P.X., J.E.V.E., R.A.G., L.M.D.D participated in manuscript preparation and finalization.

## Competing interests

The authors declare no competing financial interests. Correspondence and requests for materials should be addressed to L.M.D.D.

## Supplemental information

Supplemental information includes 5 figures and 4 tables.

## Methods

### Human myocardial samples

The right atrial appendage samples used in this study were obtained from non-diabetic and type 2 diabetic patients (aged 48-80 years) undergoing elective coronary surgery or aortic valve/arch surgery either at the Royal Melbourne Hospital, Australia, or Dunedin Hospital, New Zealand. Information on sample size is provided in the figure legends, and patient characteristics are summarized in Table S1. Tissue homogenate for glycogen and protein assays was available for 41 patients, RNA was available for 23 patients. The study was approved by the Melbourne Health Human Research and Ethics Committee and the Human and Disability Ethics Committee of New Zealand (LRS/12/01/001), and all patients provided informed consent for study participation. Right atrial appendage samples were obtained at the time of cardiac surgery and were immediately snap-frozen in liquid nitrogen and stored at −80°C for later biochemical analysis.

### Animals

Animal experiments were performed at the University of Auckland, the University of Otago, the University of Melbourne, and Cedars-Sinai Medical Center. All animal experiments were approved by the relevant institutional Animal Ethics Committee and complied with the guidelines and regulations of the Code of Practice for the Care and Use of Animals for Scientific Purposes. Animals were randomly assigned to experimental groups and were group housed (≥ 2 animals per cage) in a temperature controlled environment 21-23 °C with 12 hour light/dark cycles. At the start of all experiments, all animals were drug/test naïve. Unless otherwise specified, animals were fed regular chow. Age-matched male animals were used in all animal experiments.

#### Type 2 diabetic Zucker Diabetic Fatty (ZDF) rats

Male type 2 diabetic ZDF (*fa/fa*) rats and their non-diabetic littermates (wildtype) were sourced from Charles River, maintained on Purina 5008 diet (LabDiet) and evaluated at 20 weeks of age. ZDF rats are a well-established model of type 2 diabetes induced by a homozygous missense mutation (fatty, fa) in the leptin receptor gene (*Lepr*). ZDF rats develop type 2 diabetes from 7 weeks of age characterized by hyperinsulinemia, hyperglycemia and increased body weight^65^.

#### Type 2 diabetic db/db mice

Male db/db mice (BKS.Cg-Dock7^m^ +/+ Lepr^db^/J) and heterozygote controls (db^+^) were sourced from Charles River and evaluated at 10 weeks of age. Diabetes is the result of a point mutation in the leptin receptor gene (*Lepr, db*). db/db mice exhibit hyperglycemia, obesity, and hyperinsulinemia by 6 weeks of age^66^.

#### Type 2 diabetic high fat diet mice and rats

Obesity and type 2 diabetes was induced in male Sprague Dawley rats and C57Bl/6J mice by a high fat dietary intervention commencing at 8-9 weeks of age. After a 1 week transitional feeding period, animals were fed a high fat diet (43% kcal from fat, SF04-001, Specialty Feeds) or control reference diet (mice: 16% kcal from fat, custom mouse AIN93G control diet, Specialty Feeds; or rats: 18% kcal from fat, regular chow, Harlan, USA) for 14-15 weeks. High fat-fed mice and rats exhibit obesity, mild hyperglycemia and glucose intolerance.

#### Type 1 diabetic streptozotocin-treated rats

Type 1 diabetes was induced in male Sprague Dawley rats at 8 weeks of age via a single 55mg/kg tail vein injection of streptozotocin (STZ; Sigma), and monitored for 8 weeks. STZ-treated rats exhibit hyperglycemia from 1-2 days post-STZ injection (>16 mM), maintained throughout the 8 week diabetic period.

#### Type 2 diabetic high fat diet mice with AAV-Gabarapl1 overexpression

To evaluate the effect of cardiac-specific overexpression of *Gabarapl1* in diabetic heart disease, high fat- and control-fed male C57Bl/6J mice were randomized to treatment group after 14 weeks of the dietary intervention and received a single tail vein injection of 10^12^ gc/mouse AAV9-cTnT-*Gabarapl1* or AAV9-cTnT-Null virus (Vector Biolabs). The mice continued the high fat or control diet for a further 14 weeks (glycogen analysis) or 28 weeks (cardiomyocyte stiffness analysis).

#### Crispr-Cas9 Gabarapl1-KO mice

CRISPR-Cas9 *Gabarapl1*-knockout mice were generated by the Melbourne Advanced Genome Editing Centre using gRNAs specified in Figure S4b. C57Bl/6J embryos were injected with RNA encoding Cas9 and the specified gRNAs and inserted into pseudo-pregnant mothers. Founder mice were identified by Sanger sequencing and were crossed with C57Bl/6J mice for 2 generations. Mice used in these studies were selected from 3-7 independent founder lines.

#### Glycophagy flux assessment in type 1 diabetic rats

In a sub-cohort of animals, STZ (8 weeks post-injection) and control rats were injected with the lysosomal inhibitor, chloroquine (50 mg/kg, Sigma) or saline vehicle i.p., 4 hours prior to tissue collection, for analysis of glycophagy flux.

### Cell Culture

#### Primary neonatal rat ventricular myocytes (NRVMs)

Cardiomyocytes from male and female postnatal day 1-2 Sprague Dawley rat hearts were isolated and cultured as previously described^31^. Ventricular myocytes were isolated by enzymatic dissociation with trypsin and collagenase. Isolated cells were centrifuged (10 minutes, 1000 g), and resuspended in minimum essential media (MEM, Life Technologies) supplemented with 10 % newborn calf serum (NCBS), essential and non-essential amino acids, antibiotic-antimycotic reagent, 2 mM L-glutamine, and MEM vitamins (Life Technologies) and 26 mM NaHCO_3_ and 98 µM bromo-deoxyuridine (Sigma), and pre-plated for 1.5 hours at 37 °C to obtain a cardiomyocyte-rich fraction. Cells were plated at a cell density of 1250 cells/mm^2^ and incubated at 37 °C with 5 % CO_2_. For *Gabarapl1* gene knockdown studies, following isolation NRVMs were cultured in the MEM-NCBS media for 24 hours, then switched to antibiotic-free, serum-free Opti-MEM (Thermo Fisher Scientific) media with 100 nM siRNA (rat si*Gabarapl1*; siGENOME SMARTpool, Dharmacon; Figure S4a) vs non-targeting scrambled siRNA with Lipofectamine RNAiMAX (Life Technologies) for 16 hours. Following transfection, NRVMs were cultured for 24 hours in the MEM-NBCS media with antibiotics prior to 24 hours serum-free Dulbecco’s modified essential media (DMEM, Sigma) supplemented with 44 mM NaHCO_3_, 50 mM KCl, phenol red, 98 µM bromo-deoxyuridine, 123 µM apo-transferrin, and 1 nM insulin (Sigma), and essential and non-essential amino acids, antibiotic-antimycotic reagent, MEM vitamins, and 1 mM Na pyruvate (Life Technologies) and either control glucose (Ctrl: 5 mM glucose, 25 mM mannitol) or high glucose (HG: 30 mM glucose).

For *Gabarapl1* overexpression studies, following isolation NRVMs were cultured in MEM-NBCS as described above, for 24 hours, then switched to antibiotic-free, serum-free Opti-MEM (Thermo Fisher Scientific) with AAV9-cTnT-*Gabarapl1* or AAV9-cTnT-Null virus (10,000 gc/cell; Vector Biolabs) for 24 hours. Following transduction, NRVMs were cultured for 48 hours in standard DMEM (with supplements as described above, and 5 mM glucose) prior to 24 hours experimental conditions of control glucose (Ctrl: 5 mM glucose, 25 mM mannitol) vs high glucose (HG: 30 mM glucose). At the completion of the experimental period, cells were lysed with RIPA buffer (Thermo Fisher Scientific) for molecular analysis.

#### Murine atria-derived HL1 cardiomyocyte cell line

A stable HL1 cardiomyocyte cell line expressing GFP-tagged Gabarapl1 was established using self-inactivating retroviral particles. Non-replicative Moloney’s murine leukemia virus particles encoding GFP-Gabarapl1 were mixed with the transfection reagent, 5 µg/mL polybrene (Sigma), added to naïve HL1 cells seeded at 1×10^5^ per well, and centrifuged at 1500 g, 32 °C for 80 minutes. The transduced cells were selected by Blasticidin for 6 days. HL1-GFP-Gabarapl1 cells were maintained in Claycomb media (Sigma) supplemented with 2 mM L-glutamine (Life Technologies), 10 % NBCS (Thermo Fisher Scientific), 0.1 mM norepinephrine (Sigma) and antiobiotic-antimyotic reagent (Life Technologies), as previously described^67, 68^. Experimental conditions were defined by culture in ‘control’ standard Claycomb (Ctrl: 20 mM glucose) or high glucose Claycomb (HG: 40 mM glucose) media for 24 hours and GFP fluorescence was imaged before and after 2 hour exposure to the lysosomal inhibitor, 100 nM bafilomycin, using an epifluorescence microscope (60x oil objective, Keyence Biorevo). The number of GFP-Gabarapl1 puncta per cell was determined using ImageJ software (NIH). Cells were evaluated as either exhibiting predominantly diffuse GFP-Gabarapl1 fluorescence or exhibiting numerous punctate GFP-Gabarapl1 structures (representing Gabarapl1 accumulation in autophagosomes), and the percentage of cells with GFP-Gabarapl1 punctate was calculated for each image as previously described^69^.

#### Human pluripotent stem cell-derived cardiomyocytes (hPSC-CMs)

Human pluripotent stem cells were maintained and differentiated into cardiomyocytes (hPSC-CMs) as previously described^70, 71^ with ethical approval from The University of Queensland’s Medical Research Ethics Committee under the regulations of the National Health and Medical Research Council of Australia (NHMRC). hPSC-CMs were dissociated and seeded onto gelatin coated coverslips at 100,000 cells/cm^2^ in α-MEM GlutaMAX (Thermo Fisher Scientific), 10 % Fetal bovine serum (FBS; Thermo Fisher Scientific), 200 μM L-ascorbic acid 2-phosphate sesquimagnesium salt hydrate (Sigma) and 1 % penicillin/streptomycin (Thermo Fisher Scientific). After 48 hours, cells were cultured in maturation medium comprising DMEM, glutamine and phenol red (Thermo Fisher Scientific) supplemented with 4 % B27 (Thermo Fisher Scientific), 1 % GlutaMAX (Thermo Fisher Scientific), 200 μM L-ascorbic acid 2-phosphate sesquimagnesium salt hydrate (Sigma) and 1 % penicillin/streptomycin (Thermo Fisher Scientific), with either control glucose (Ctrl: 1 mM glucose with 24 mM mannitol) or high glucose (HG: 25 mM glucose) conditions for 72 hours and lysed for glycogen analysis, or fixed for electron microscopy as described previously^70, 72^.

#### Human cardiac organoids (hCOs)

hCOs were generated from hPSC-CMs using Heart-Dyno culture inserts and cultured in maturation media as previously described^70^. At Day 9 hCOs were treated with AAV9-cTnT-*Gabarapl1* or AAV9-cTnT-Null virus (5×10^9^ gc/hCO; Vector Biolabs) for 72 hours in maturation media. hCOs were subsequently washed and cultured in control glucose (Ctrl: maturation media with 1 mM glucose with 24 mM mannitol) or high glucose (HG: maturation media with 25 mM glucose) conditions for 4 days. hCO contractile properties and kinetics were assessed as previously described^70^. At the completion of contraction analysis, a subset of cardiac organoids were fixed for electron microscopy as described previously^70, 72^.

### Glucose tolerance testing

Glucose tolerance testing was performed in *Gabarapl1*-KO and T2D-AAV mice following 6 hours fasting^73^. Baseline blood glucose levels were measured using an Accu-Chek glucometer with a small blood sample obtained from a needle prick to the tail vein. Glucose (1.5 g/kg body weight) was injected i.p. and blood glucose was measured at 5, 15, 30, 60 and 90 minutes after the glucose injection.

### Echocardiography

Animals were anaesthetized with isoflurane and transthoracic echocardiography was performed using the GE Vivid 9 Dimension echocardiography platform with a 15 MHz i13L linear array transducer (GE Healthcare). Left ventricular (LV) m-mode two-dimensional echocardiography was performed in parasternal short axis view to measure LV wall and chamber dimensions to derive systolic function parameters: % ejection fraction ((end diastolic volume – end systolic volume)/end diastolic volume) ×100), and % fractional shortening ((LV end diastolic diameter – LV end systolic diameter)/(LV end diastolic diameter) ×100). Pulse wave Doppler and tissue Doppler imaging were acquired from the apical 4 chamber view to assess LV diastolic function parameters: velocity of early mitral inflow (E) and early diastolic velocity of mitral annulus (e′) and E/e′ ratio. Three consecutive cardiac cycles were sampled for each measurement taken, and analysis was performed in a blinded manner.

### Adult cardiomyocyte isolation and nano-mechanical analysis

Adult rodent cardiomyocytes were isolated as previously described (mouse^74^; rat^75^), attached to glass rods (Myotak, Ionoptix) and lifted from the coverslip using motorized micromanipulators (MC1000E, Siskiyou Corporation, OR, USA). Cardiomyocytes were paced (1 Hz (rat) or 2 Hz (mouse), 2.0 mM Ca^2+^, 37 °C) and subjected to progressive longitudinal stretch (Nano-drive, Mad City Labs Inc.). Sarcomere length/shortening, force development (force transducer stiffness 20 N/m; Ionoptix) and intracellular Ca^2+^ transients (Fura2, 5 μM, microfluorimetry F_340:380nm_ ratio) were simultaneously evaluated (Myostretcher; Ionoptix). Diastolic force-length relationships were constructed by plotting diastolic force against % cardiomyocyte stretch. All indices were analyzed off-line using IonWizard software (IonOptix) and were determined after averaging 10 steady-state transients for each myocyte.

### Molecular analyses

#### Glycogen content

Glycogen content was measured by digesting an aliquot of tissue homogenate with amyloglucosidase (#10102857001, Roche) at 50 °C for 60 minutes in 1 % triton-X, 0.1 M Na acetate, pH 6.0. Following centrifugation at 16,000 g, 4 °C for 2 minutes, the supernatant was assayed for glucose concentration using glucose oxidase/peroxidase colorimetric glucose assay with 45 μg/ml o-dianisdine dihydrochloride (Sigma-Aldrich), and absorbance measured at 450 nm. Another tissue homogenate aliquot was processed in parallel without amyloglucosidase to determine background glucose content. Glycogen levels are presented as glucose units (nmol), normalized to protein (mg, determined by Lowry assay) and depicted as relative levels for comparative purposes.

#### Subcellular fractionation

Frozen tissue was homogenized in lysis buffer (100 mM Tris-HCl (pH 7.0), 5 mM EGTA, 4 mM EDTA, protease and phosphatase inhibitors (Roche)) and centrifuged at 16,000 g (10 minutes, 4 °C). The supernatant was collected for the cytosolic fraction and the pellet was resuspended in lysis buffer with 1 % triton (Sigma) and centrifuged at 16,000 g (10 minutes, 4 °C). The detergent-soluble supernatant was collected for the crude membrane fraction.

#### Immunoblotting

Sample protein concentrations were determined using a Lowry assay. Prior to immunoblotting, tissue homogenates were prepared in loading buffer (50 mM Tris-HCl (pH 6.8), 2 % sodium dodecyl sulfate, 10 % glycerol, 0.1 % bromophenol blue and 2.5 % 2- mercaptoethanol), and equal amounts of protein were loaded into the SDS-PAGE gel. Antibodies for the following proteins were used: phosphorylated (Ser641) glycogen synthase (ab81230, Abcam), glycogen synthase (3893, Cell Signaling), phosphorylated (Ser14) glycogen phosphorylase (gift from Dr David Stapleton), glycogen phosphorylase (gift from Dr David Stapleton), and LC3B (2775, Cell Signaling) and Gabarapl1 (26632, Cell Signaling). Membranes were incubated with anti-rabbit horseradish peroxidase-conjugated secondary antibody (GE Healthcare). The ECL Prime (Amersham, GE Healthcare) chemiluminescent signal was visualized with a ChemiDoc-XRS Imaging device and band intensity quantified using ImageLab software (Bio-Rad). Equal protein loading was confirmed by Coomassie staining of polyvinylidene difluoride membranes (Coomassie Brilliant Blue R-250, Bio-Rad).

#### Quantitative RT-PCR

RNA was extracted from frozen cardiac tissues using the TRIzol® reagent in conjunction with the PureLink™ Micro-to-Midi Total RNA Purification kit (Life Technologies) with on-column DNase treatment (PureLink DNase; Life Technologies). RNA was reversed transcribed as per the manufacturer’s instructions (#18080-051; Life Technologies). Real-time PCR was performed using Sybr-Green (Life Technologies) with the following primer pairs: mouse and rat *Gabarapl1*, 5’-GGTCATCGTGGAGAAGGCTC-3’ (forward) and 5’-TAGAACTGGCCAACAGTGAGG-3’ (reverse); 18S housekeeper gene, 5’-TCGAGGCCCTGTAATTGGAA-3’ (forward) and 5’-CCCTCCAATGGATCCTCGTT-3’ (reverse). The comparative ΔΔCt method was used to analyze the genes of interest as described^76^.

For assessment of viral copy number in AAV9-treated mouse heart tissue, DNA was extracted as per the manufacturer’s instructions (K0152, Thermo Fisher Scientific). AAV9- treated mouse heart DNA was subjected to digital droplet qPCR as per the manufacturer’s instructions (Biorad) using primers targeting the WPRE component of the viral vector: 5’-CTGGTTGCTGTCTCTTTATGAGGAG-3’ (forward) and 5’-CACTGTGTTTGCTGACGCAACC-3’ (reverse). Viral copies/ μg DNA was quantified using QuantaSoft^TM^ Analysis Pro.

#### Glycogen proteomics

Cardiac (left ventricle) and skeletal muscle (quadriceps) tissue was collected from control and glycemic challenged Sprague Dawley rats (2 days post-destruction of pancreatic β-cells with 55mg/kg STZ tail vein injection, a time-point when blood glucose levels were >50% elevated from baseline). Glycogen was extracted as previously described^52^. Briefly, tissues were homogenized in glycogen extraction buffer (in mM: 50 Tris pH 8, 150 NaCl, 2 EDTA) and cellular debris was removed by centrifugation (6000 g, 10 minutes, 4 °C). The supernatant was ultracentrifuged (300,000 g, 60 minutes, 4 °C), and the resulting pellet was resuspended in glycogen extraction buffer and layered over a sucrose gradient (75, 50, 25 % w/v sucrose) and ultracentrifuged (300,000 g, 2 hours, 4 °C). The resulting pellet containing glycogen was resuspended in glycogen extraction buffer. Glycogen-associated proteins were displaced from the glycogen by addition of α1,4 malto-oligosaccharides (maltodextrin, 50 mg/ml) and ultracentrifuged (400,000 g, 20 minutes, 4 °C). The supernatant containing proteins eluted from the glycogen were reduced (tris(2-carboxyethyl)phosphine (TCEP), 5 mM), alkylated (iodoacetamide, 10 mM), trypsin-digested (1:75 w/w, Promega) and desalted (HLB μElution plate, Waters) in preparation for liquid chromatography tandem mass spectrometry (LC-MS/MS). LC-MS/MS was performed in micro-flow mode (Eksigent 415 LC, 5600+ TripleTOF Sciex) using a trap column (ChromXP C18CL 10×0.3 mm 5 µm 120 Å; flow rate 10 µL/min, 3 minutes), and an analytical column (ChromXP C18CL 150×0.3 mm 3 µm 120 Å; flow rate 5 µL/min, 30 °C). Peptides were eluted in a linear gradient spanning 3-35 % acetonitrile in water (0.1 % Formic Acid) over 60 minutes. MS/MS scans were acquired as previously described^77^. All raw data from the 5600+ TripleTOF mass spectrometer were converted to mzXML using a Sciex Data converter. Data were analyzed using the Sorcerer 2TM-SEQUEST® algorithm (Sage-N Research, Milpitas CA, USA) searched against the concatenated target/decoy Rat Uniprot 2018 reviewed FASTA database^78, 79^ limited to trypsin-digested peptides, 50 PPM parent ion tolerance, 1 Da fragment ion tolerance, carbamidomethyl of cysteine fixed modification and oxidation of methionine variable modification. Quantitation, validation and normalization was performed using Scaffold 4 on proteins defined by a minimum of 2 proteotypic peptides (5 % false discovery rate) with >99.0 % probability (Protein Prophet algorithm^80^). Relative protein quantification was derived from MS/MS data using spectral counting, subjected to normalized spectral abundance factor (NSAF)^81^ and a minimum value of 0 assigned to non-detected proteins. A subset of proteins identified to be i) exclusively detected in control samples, ii) exclusively detected in glycemic challenge samples and iii) differentially abundant in glycemic challenge vs control samples (-0.2 ≤ fold change ((STZ/control)-1) ≥ 0.2, p<0.05) were used for subsequent analyses. Proteins relating to the contractile apparatus and protein translation GO categories were excluded. Venn diagrams were constructed for cardiac and skeletal muscle datasets to show the number of unique or differentially abundant proteins in control vs 48 hour glycemic challenge samples identified from the Scaffold 4 output. Proteins uniquely detected in the glycemic challenge samples (not detected in control samples) were categorized using Enrichr software^80, 82^ into GO categories Carbohydrate Metabolic Processes (GO0005975), Fatty Acid Metabolic Processes (GO0006629) and Amino Acid Metabolic Processes (GO0006520) and Log_2_ protein abundance presented in a Bubble Chart format. GO analysis (Enrichr) of all uniquely detected and differentially abundant proteins identified the top 7 GO Biological Processes categories associated with the glycogen proteome response to glycemic challenge for cardiac and skeletal muscle tissues. The normalized spectral counts of the uniquely detected and differentially abundant proteins in the GO category Carbohydrate Metabolic Processes were converted to heatmap intensity for graphical representation. The MS/MS proteomics data have been deposited to the ProteomeXchange Consortium (http://www.proteomexchange.org)83 via the PRIDE partner repository with the dataset identifier #PXD009658.

#### STRING functional network analysis

The glycogen-tagging protein, Stbd1, identified to be of importance in the glycogen proteome response to glycemic challenge, was subjected to STRING protein-protein association network analysis V11.0^84^ to identify common pathways or associations for subsequent investigation. Each association identified was based on text-mining and/or experimentally determined evidence in the rat database. The minimum accepted confidence score (minimum probability) for an association was set at 0.7. Primary (max. 10) and secondary (max. 20) interactions were identified and presented as a network graph.

#### Gene profiling and Jensen DISEASE classification

Gene profiling was performed on RNA extracted from type 1 diabetic rat heart tissue (STZ-treated), reversed transcribed to cDNA using the RT^2^ First Strand kit (Qiagen). RNA and cDNA quality was verified using the RT^2^ RNA Quality Control PCR Array (Qiagen). Custom PCR array plates were designed with 57 genes of interest and housekeeping gene β-actin (Table S1). Real time PCR was performed and differential gene expression was determined by ΔΔCt analysis. Differentially expressed genes were subjected to disease classification using the Jensen DISEASE resource (http://diseases.jensenlab.org/). The top 10 diseases most closely aligned with the differentially detected genes in T1D hearts were presented relative to their combined score calculated as ln(p-value) x odds ratio.

### Electron microscopy

Left ventricular free wall heart segments (1 mm^3^) were fixed in 2.5 % glutaraldehyde in 0.1 M phosphate buffer for >24 hours at 4 °C. Fixed hearts were processed in 1 % osmium tetroxide/1.5 % potassium ferrocyanide in 0.1 M phosphate buffer for 90 minutes and dehydrated with increasing ethanol concentration followed by propylene oxide (PO; Merck-Millipore). Samples were infiltrated and embedded in pure resin at 60 °C for >48 hours. Resin blocks were cut (1 μm) and stained with 1 % toluidine blue in 1 % Na tetraborate prior to cutting of 80 nm sections on copper EM grids and staining with uranyl acetate for 30 minutes and 0.3 % lead citrate for 3 minutes. Sections were imaged systematically through the grid locations using a Tecnai™ G² Spirit Twin transmission electron microscope.

### Histological analysis

#### Periodic-acid Schiff glycogen staining

Heart mid-sections were fixed in 10 % formalin (Sigma) for 24 hours, embedded in paraffin and cut into 4 μm sections. Following dewaxing, sections were oxidized with 1 % periodic acid for 5 minutes, washed and incubated in Schiff’s reagent for 10 minutes at room temperature. Sections were then washed and counterstained with Mayer’s haematoxylin for 30 seconds, washed, dehydrated and mounted in DPX. Selected sections were incubated with 5 mg/ml α-amylase for 5 minutes at 37 °C (Sigma) to confirm distinct glycogen and glycoprotein stain localization. Images were acquired using a Zeiss Imager DI microscope with a Zeiss AxioCam MRc5 colour camera (Carl Zeiss).

#### Picrosirius Red collagen staining

Heart mid-sections were fixed in 10 % formalin (Sigma) for 24 hours, embedded in paraffin and cut into 6 μm sections. Sections were stained with 0.1 % Sirius red in 1.2 % w/v picric acid for 1 hour, washed and dehydrated. Sections were scanned with a brightfield digital scanner using (Panoramic ScanII Scanner, 3D Histech) and images captured using CaseViewer version 2.2 acquisition software (3DHistech; 20x magnification).

### Statistics and reproducibility

Data are presented as mean ± SEM (with the exception of data derived from clinical samples presented as mean ± SD) and statistical analysis was performed using Graphpad Prism V7.0. A description of biological replicates is provided within the figure legends. All datasets were tested for normal distribution using Shapiro-Wilk tests and for equal variances using F-tests (2 groups) or Bartlett’s test (2 independent variables). For comparison between two groups, a 2-sided Student’s t-test was used. For assessment between two independent variables, two-way ANOVA with Bonferroni multiple comparisons post-hoc test was used. Some datasets required logarithmic transformation to meet equal variance and/or normal distribution statistical assumptions for parametric testing. In datasets where transformation did not achieve equal variance and normal distribution, non-parametric tests were used (2 groups: Mann Whitney U; 2 variables: Kruskal-Willis). For correlation analyses Pearson’s correlation coefficient were used. A p-value of <0.05 was considered statistically significant.

## Data availability

The datasets generated during and/or analyzed during the current study are available from the corresponding author on reasonable request.

## Notes

### Competing Interest Statement

The authors have declared no competing interest.

