## Supplementary tables & figures for "Protective role of the Atg8 homologue Gabarapl1 in regulating cardiomyocyte glycophagy in diabetic heart disease"

**Table S1. Characteristics of non-diabetic and type 2 diabetic patients.** Related to Figure 1a & 3d.

| | Non-diabetic | Type 2 diabetic |
|---|---|---|
| n (total) | 26 | 21 |
| n (glycogen) | 23 | 18 |
| n (PCR) | 12 | 11 |
| Age (years) | 67.8 ± 6.3 | 66.5 ± 9.1 |
| Sex | 4F + 22M | 6F + 15M |
| Beta blockers | 14/26 | 18/21 |
| ACE inhibitors/Angiotensin Receptor Blockers | 18/26 | 16/21 |
| Statins | 19/26 | 19/21 |
| Metformin | 0/26 | 15/21 |
| Insulin | 0/26 | 9/21 |
| Hypertension | 16/26 | 17/21 |

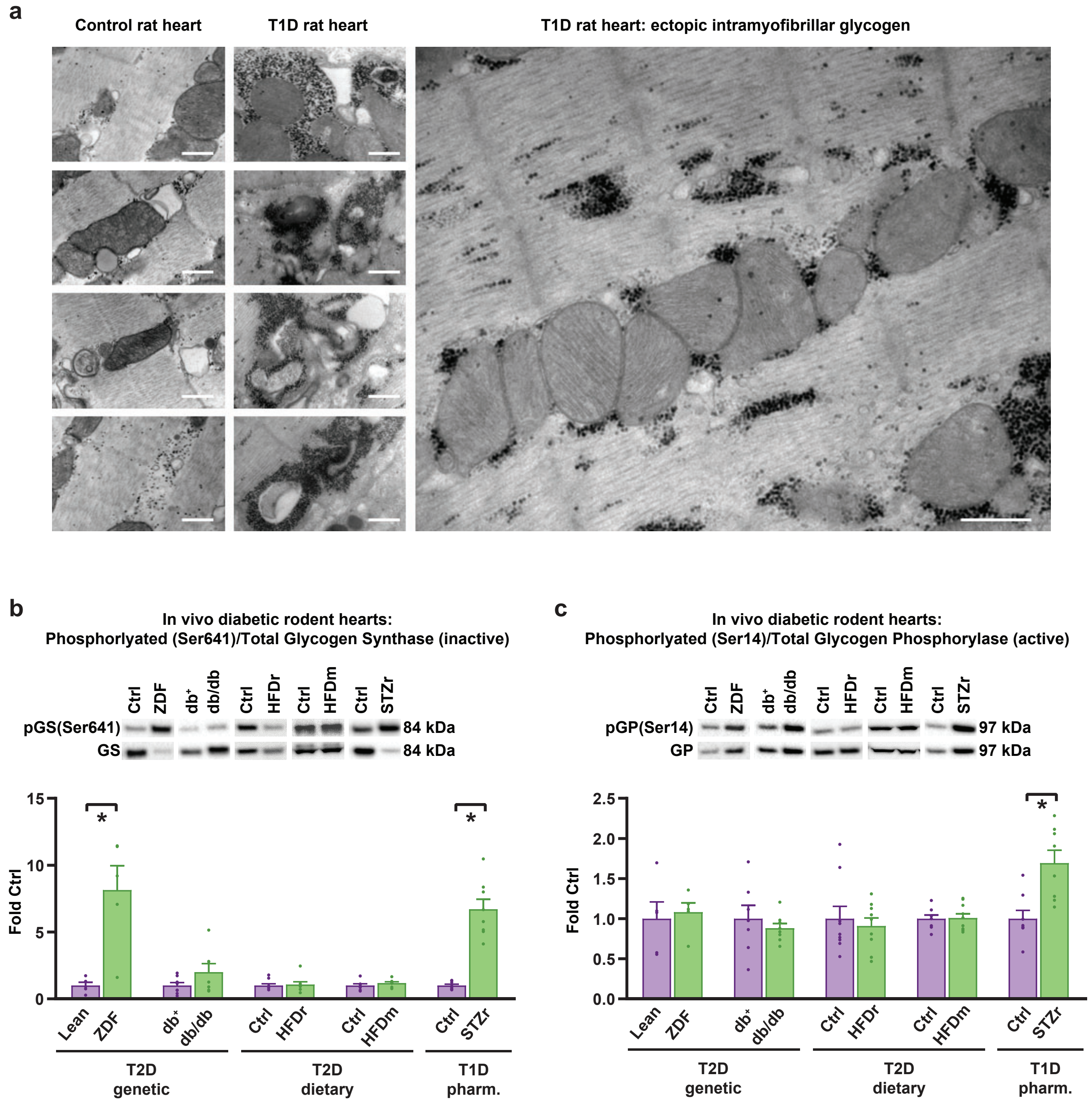


**Figure S1. Transmission electron micrographs of type 1 diabetic myocardium and cardiac glycogen synthase and phosphorylase expression in diabetic rodent models. a,** Transmission electron microscopy images of control (left) and type 1 diabetic (middle & right) rat heart sections (T1D, streptozotocin-induced). Electron-dense glycogen is evident as black particles, scale bar 500nm. **b,** Glycogen synthase activity is inhibited (increased phosphorylation at Ser641) in ZDF and STZ rat diabetic hearts and unchanged in db/db, HFD rat and HFD mouse hearts (n ≥ 5). **c,** Glycogen phosphorylase activity is increased (increased phosphorylation at Ser14) in STZ rat hearts and unchanged in ZDF, db/db, HFD rat and HFD mouse hearts (n ≥ 5). Data are presented as mean ± s.e.m. *p<0.05. Related to Figure 1a & 1b.

**Table S2. List of genes analyzed by PCR array in myocardial tissue from type 1 diabetic rats.** Related to Figure 1d.

| ACCESSION# | GENE NAME | GENE ID |
|---|---|---|
| NM_001012097 | Autophagy Related 7 | Atg7 |
| NM_001013988 | Starch Binding Domain 1 | Stbd1 |
| NM_001014152 | Phosphorylase Kinase Beta Subunit | Phkb |
| NM_001014218 | Autophagy Related 9a | Atg9a |
| NM_001014250 | Autophagy Related 5 | Atg5 |
| NM_001024743 | UDP-Glucose Pyrophosphorylase 2 | Ugp2 |
| NM_001025018 | DNA-Damage Regulated Autophagy Modulator 2 | Dram2 |
| NM_001025711 | Autophagy Related 4b, Cysteine Peptidase | Atg4b |
| NM_001038495 | Autophagy Related 12 | Atg12 |
| NM_001044294 | Gaba Type A Receptor Associated Protein Like 1 | Gabarapl1 |
| NM_001077675 | Tripartite Motif Containing 72 | Trim72 |
| NM_001106395 | Forkhead Box O3 | FoxO3 |
| NM_001106694 | PTEN Induced Putative Kinase 1 | Pink1 |
| NM_001106723 | Phosphatidylinositol-4,5-Bisphosphate 3-Kinase, Catalytic Subunit Gamma | Pik3cg |
| NM_001107335 | Death Associated Protein Kinase 1 | Dapk1 |
| NM_001107627 | Sirtuin 1 | Sirt1 |
| NM_001107948 | Autophagy Related 4c, Cysteine Peptidase | Atg4c |
| NM_001108341 | Unc-51 Like Autophagy Activating Kinase 1 | Ulk1 |
| NM_001108564 | Amylo-Alpha-1, 6-Glucosidase, 4-Alpha-Glucanotransferase | Agl |
| NM_001108777 | Phosphoinositide-3-Kinase, Regulatory Subunit 4 | Pik3r4 |
| NM_001108809 | Autophagy Related 16-Like 1 | Atg16l1 |
| NM_001109615 | Glycogen Synthase 1 | Gys1 |
| NM_001127297 | WD Repeat Domain, Phosphoinositide Interacting 1 | Wipi1 |
| NM_001134341 | Autophagy And Beclin 1 Regulator 1 | Ambra1 |
| NM_001191560 | Autophagy Related 16-Like 2 | Atg16l2 |
| NM_001191846 | Forkhead Box O1 | FoxO1 |
| NM_001276762 | Epilepsy, Progressive Myoclonus Type 2A | Epm2a |
| NM_012565 | Glucokinase | Gck |
| NM_012638 | Phosphorylase, Glycogen, Muscle | Pygm |
| NM_012735 | Hexokinase 2 | Hk2 |
| NM_012751 | Solute Carrier Family 2 Member 4 | GLUT4 |
| NM_012857 | Lysosomal-Associated Membrane Protein 1 | Lamp1 |
| NM_012922 | Caspase 3 | Casp3 |
| NM_016993 | B-Cell Cll/Lymphoma 2 | Bcl2 |
| NM_017068.2 | Lysosomal-Associated Membrane Protein 2 | Lamp2 |
| NM_017320 | Cathepsin s | Ctss |
| NM_017344 | Glycogen Synthase Kinase 3 Alpha | Gsk3a |
| NM_019142 | Protein Kinase Amp-Activated Catalytic Subunit Alpha 1 | Prkaa1 |
| NM_019906 | Mechanistic Target Of Rapamycin | Mtor |
| NM_022597 | Cathepsin b | Ctsb |
| NM_022626 | Phosphorylase Kinase, Alpha 1 | Phka1 |
| NM_022698 | Bcl2-Associated Agonist Of Cell Death | Bad |
| NM_022706 | Gaba Type A Receptor Associated Protein Like 2 | Gabarapl2 |
| NM_022867 | Microtubule-Associated Protein 1 Light Chain 3 Beta | Map1lc3b |
| NM_022958 | Phosphatidylinositol 3-Kinase, Catalytic Subunit Type 3 | Pik3c3 |
| NM_031573 | Phosphorylase Kinase, Gamma 1 | Phkg1 |
| NM_031715 | Phosphofructokinase, Muscle | Pfkm |
| NM_032080 | Glycogen Synthase Kinase 3 Beta | Gsk3b |
| NM_033230 | Akt Serine/Threonine Kinase 1 | Akt1 |
| NM_053420 | Bcl2/Adenovirus E1B Interacting Protein 3 | Bnip3 |
| NM_053494 | Solute Carrier Family 2 Member 8 | GLUT8 |
| NM_053739 | Beclin 1 | Becn1 |
| NM_134394 | Autophagy Related 3 | Atg3 |
| NM_138827 | Solute Carrier Family 2 Member 1 | GLUT1 |
| NM_172036 | Gaba Type A Receptor-Associated Protein | Gabarap |
| NM_178866 | Insulin-Like Growth Factor 1 | Igf1 |
| NM_181550 | Sequestosome 1 | Sqstm1 |
| NM_199118 | Glucosidase, Alpha, Acid | Gaa |
| NM_199500 | Microtubule-Associated Protein 1 Light Chain 3 Alpha | Map1lc3a |

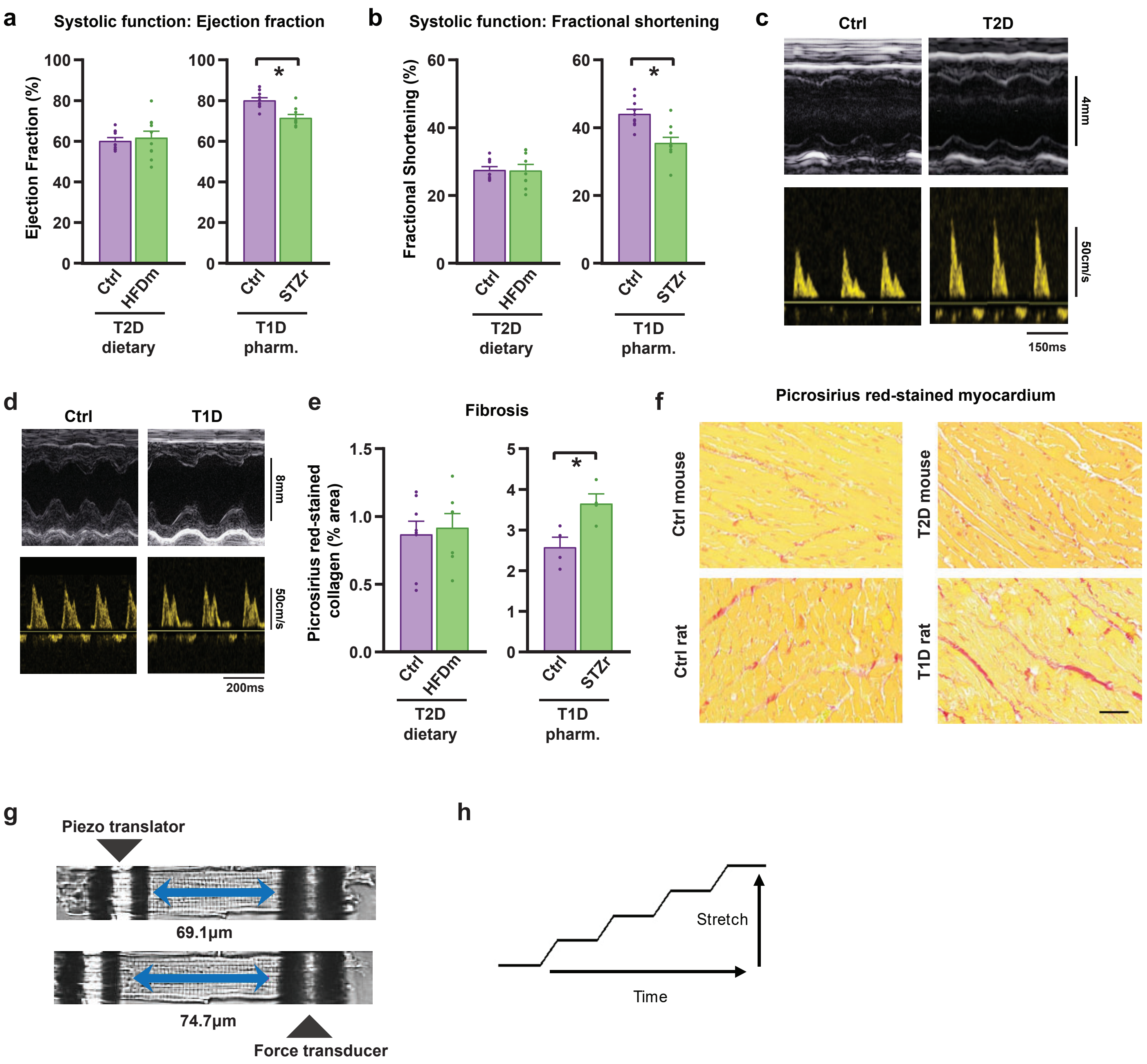


**Figure S2. Cardiac systolic functional parameters and quantification of fibrosis in diabetic rodents.** Echocardiography M-mode-derived ejection fraction **a,** and fractional shortening **b,** are unchanged with diabetes in type 2 diabetic mice (14 week high fat diet (HFD), n≥9 animals) and decreased in type 1 diabetic rats (streptozotocin (STZ)-induced, n ≥9 animals). **c,** Exemplar M-mode and flow Doppler echocardiography traces in type 2 diabetic mice (HFD). **d,** Exemplar M-mode and flow Doppler echocardiography traces in type 1 diabetic rats (STZ). **e,** Cardiac fibrosis is unchanged in type 2 diabetic mouse hearts (14 week high fat diet, n≥7 animals) and increased in type 1 diabetic rat hearts (STZ-induced, n=4 animals), evidenced by histological quantification of picrosirius red-stained collagen in paraffin-embedded formalin-fixed sections. **f,** Exemplar images of picrosirius red-stained myocardial sections from type 2 diabetic mice (HFD) and type 1 diabetic rats (STZ). Scale bar, 50µm. **g,** Exemplar images of a non stretched and stretched cardiomyocyte attached between two glass rods. **h,** Cardiomyocyte stretch protocol. Data are presented as mean ± s.e.m. *p<0.05. Related to Figures 1e-h.

**Table S3. Proteomic (LC-MS/MS) analysis of glycogen-associated proteins in cardiac and skeletal muscle in control and 48hr glycemic challenge rats.** Differentially abundant and uniquely detected proteins from the Carbohydrate Metabolic Processes GO category in the glycogen proteome of cardiac and skeletal muscle from glycemic challenge rats. In vivo 48hr glycemic challenge was induced by 55mg/kg i.p. streptozotocin destruction of pancreatic b-cells (n=4 rats/group). Related to Figure 2d.

| ACCESSION# | PROTEIN NAME | GENE NAME |
|---|---|---|
| ALAT1_RAT | Alanine aminotransferase 1 | Gpt |
| CISY_RAT | Citrate synthase, mitochondrial | Cs |
| CLH1_RAT | Clathrin heavy chain 1 | Cltc |
| DHSO_RAT | Sorbitol dehydrogenase | Sord |
| ENOG_RAT | Gamma-enolase | Eno2 |
| F16P2_RAT | Fructose-1,6-bisphosphatase isozyme 2 | Fbp2 |
| GSTO1_RAT | Glutathione S-transferase omega-1 | Gsto1 |
| HXK1_RAT | Hexokinase-1 | Hk1 |
| HXK2_RAT | Hexokinase-2 | Hk2 |
| IDHC_RAT | Isocitrate dehydrogenase [NADP] cytoplasmic | Idh1 |
| KPB1_RAT | Phosphorylase b kinase regulatory subunit alpha, skeletal muscle isoform | Phka1 |
| LGUL_RAT | Lactoylglutathione lyase | Glo1 |
| MA2C1_RAT | Alpha-mannosidase 2C1 | Man2c1 |
| MDHC_RAT | Malate dehydrogenase, cytoplasmic | Mdh1 |
| MTOR_RAT | Serine/threonine-protein kinase mTOR | Mtor |
| PFKAL_RAT | ATP-dependent 6-phosphofructokinase, liver type | Pfkl |
| PFKAP_RAT | ATP-dependent 6-phosphofructokinase, platelet type | Pfkp |
| PGAM1_RAT | Phosphoglycerate mutase 1 | Pgam1 |
| PGK1_RAT | Phosphoglycerate kinase 1 | Pgk1 |
| PHKG1_RAT | Phosphorylase b kinase gamma catalytic chain, skeletal muscle/heart isoform | Phkg1 |
| PK3CA_RAT | Phosphatidylinositol 4,5-bisphosphate 3-kinase catalytic subunit alpha isoform | Pik3ca |
| PP1B_RAT | Serine/threonine-protein phosphatase PP1-beta catalytic subunit | Ppp1cb |
| STBD1_RAT | Starch-binding domain-containing protein 1 | Stbd1 |
| TPIS_RAT | Triosephosphate isomerase | Tpi1 |

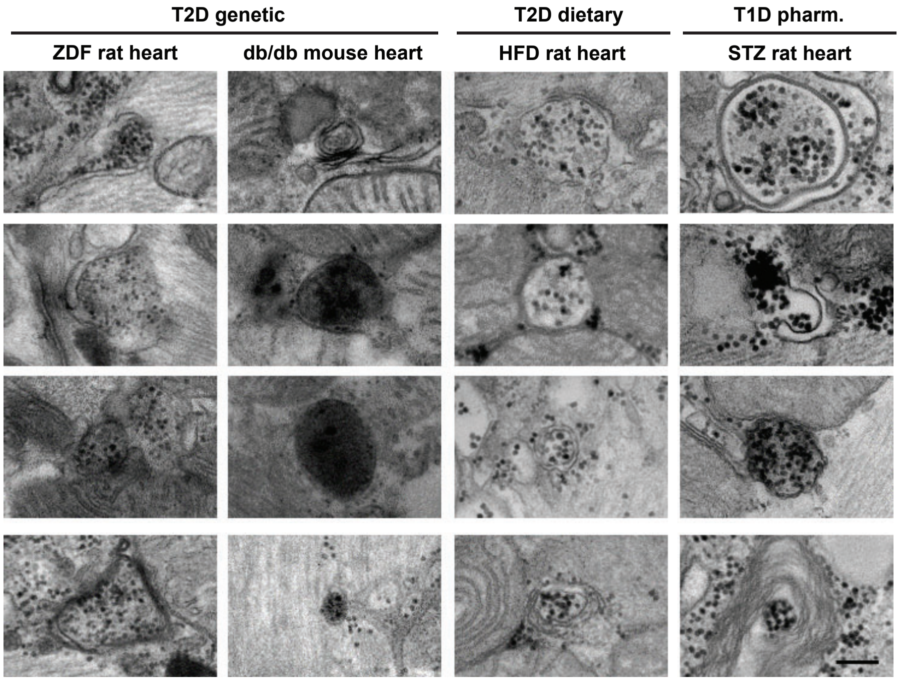


**Figure S3. Transmission electron micrographs of glycogen-filled autophagosomes (glycophagosomes) in diabetic myocardium.** Transmission electron microscopy images of diabetic rat heart sections containing glyco-phagosomes. Electron-dense glycogen is evident as black particles, scale bar 200nm. T2D, type 2 diabetes; T1D, type 1 diabetes; ZDF, Zucker diabetic fatty rat 20wk old; dbdb, 10wk old; HFD, 14wk high fat diet; STZ, 8wk streptozotocin-treated. Related to Figure 2g.

**a** **siRNA sequences: *Gabarapl1* knockdown in vitro**

| | siRNA sequence (5'-3') | Genic location |
|---|---|---|
| Seq1 | UUCCCGUAGACACUUUCAU | Exon4: protein coding |
| Seq2 | AUUGCGAACAGCCCUAUUU | Exon4: UTR |
| Seq3 | UUUAACGCCAUCCAAACUG | Exon4: UTR |
| Seq4 | UCGCUUUGCAUUCCAGUGC | Exon4: UTR |

**b** **Crispr-Cas9 sequences: *Gabarapl1*-KO in vivo**

***Gabarapl1*-KO sequence**

Chr 6: 129,536,667 – 129,536,701 | 129,542,753 – 129,542,789

| | |
|---|---|
| Predicted | ATGCTCTTAGATGGCCCCAGTCTTTGGCCTGGTG - ACTGGCAGCCATGTAGGCAGTTCACCATGAGTAGGT |
| Line 1 | ATGCTCTTAGATGGCCCCAGTCTTTGGCCTGG- - - - - - - - CAGCCATGTAGGCAGTTCACCATGAGTAGGT |
| Line 2 | ATGCTCTTAGATGGCCCCAGTCTTTGGCCTGGTGGACTGGCAGCCATGTAGGCAGTTCACCATGAGTAGGT |
| Line 3 | ATGCTCTTAGATGGCCCCAGTCTTTGGCCTGG- - - - - - - - CAGCCATGTAGGCAGTTCACCATGAGTAGGT |
| Line 4 | ATGCTCTTAGATGGCCCCAGTCTTTGGACTGG- - - - - - - - CAGCCATGTAGGCAGTTCACCATGAGTAGGT |
| Line 5 | ATGCTCTTAGATGGCCCCAGTCTTTGGCCTGG- - - - - - - - CAGCCATGTAGGCAGTTCACCATGAGTAGGT |
| Line 6 | ATGCTCTTAGATGGCCCCAGTCTTTGGACTGG- - - - - - - - CAGCCATGTAGGCAGTTCACCATGAGTAGGT |
| Line 7 | ATGCTCTTAGATGGCCCCAGTCTTTGGCCTGG- - - - - - - - CAGCCATGTAGGCAGTTCACCATGAGTAGGT |

**guideRNA sequences: *Gabarapl1* CRISPR knockout**

| | Spacer sequence (5'-3') | Strand | PAM (5'-3') | Genic location | Genomic location |
|---|---|---|---|---|---|
| gRNA1 | ggtggtgcgtcaaactatcg | + | cctggtg | Intron | Chr6:129,536,697 |
| gRNA2 | ggtctggtcccagatttgac | + | actggc | post-gene | Chr6:129,542,735 |

**c** ***Gabarapl1*-KO in vivo validation**

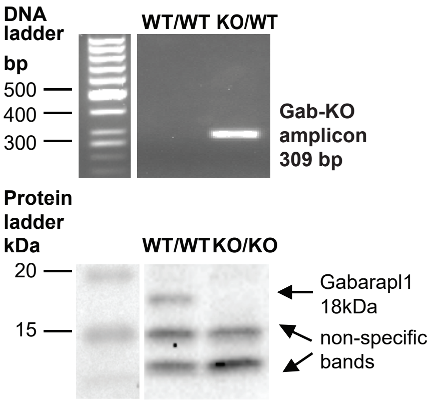


**d** ***Gabarapl1*-KO (30wk): *Gabarapl1* mRNA**

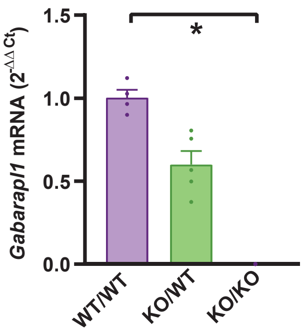


**e** **Characteristics of 12 week old *Gabarapl1*-KO mice.**

| | WT/WT | WT/KO |
|---|---|---|
| Age (weeks) | 12.8 ± 0.2 | 12.5 ± 0.2 |
| Heart weight (mg) | 131.0 ± 4.6 | 137.5 ± 2.2 |
| Heart weight/body weight (mg/g) | 4.7 ± 0.1 | 4.8 ± 0.1 |
| Tibia length (mm) | 16.9 ± 0.2 | 16.9 ± 0.2 |
| Heart weight/tibia length (mg/mm) | 7.8 ± 0.3 | 8.2 ± 0.2 |

**Figure S4. *Gabarapl1* knockdown characteristics *in vitro* (siRNA) and *in vivo* (Crispr Cas9 mice). a,** Details of the siRNA sequences used to transfect neonatal rat ventricular myocytes to knockdown *Gabarapl1* gene expression. **b,** *Gabarapl1* sequence details for the 7 founder Crispr-Cas9 mouse lines confirming excision of exons 2-4 from the *Gabarapl1* gene (Next Generation sequencing). Text in red highlights mutations differing from the predicted sequence. **c,** Validation of *Gabarapl1* knockdown using DNA electrophoresis to identify the presence of the *Gabarapl1*-KO allele in the heterozygote *Gabarapl1*-KO (tail sample), and immunoblot to confirm the absence of the Gabarapl1 band in the homozygote knockout mouse (membrane-enriched fraction of heart homogenate). **d,** Heterozygote *Gabarapl1*-KO mice exhibit ~50% knockdown of the *Gabarapl1* gene (qPCR) in the heart, and absence of the *Gabarapl1* gene is confirmed in the homozygote *Gabarapl1*-KO mouse heart (30 week old male mice, n=4-5 mice). **e,** Characteristics of 12 week old male heterozygote *Gabarapl1*-KO mice (n=8-12 mice). Data are presented as mean ± s.e.m. *p<0.05. Related to Figures 3f-h and 4a-i.

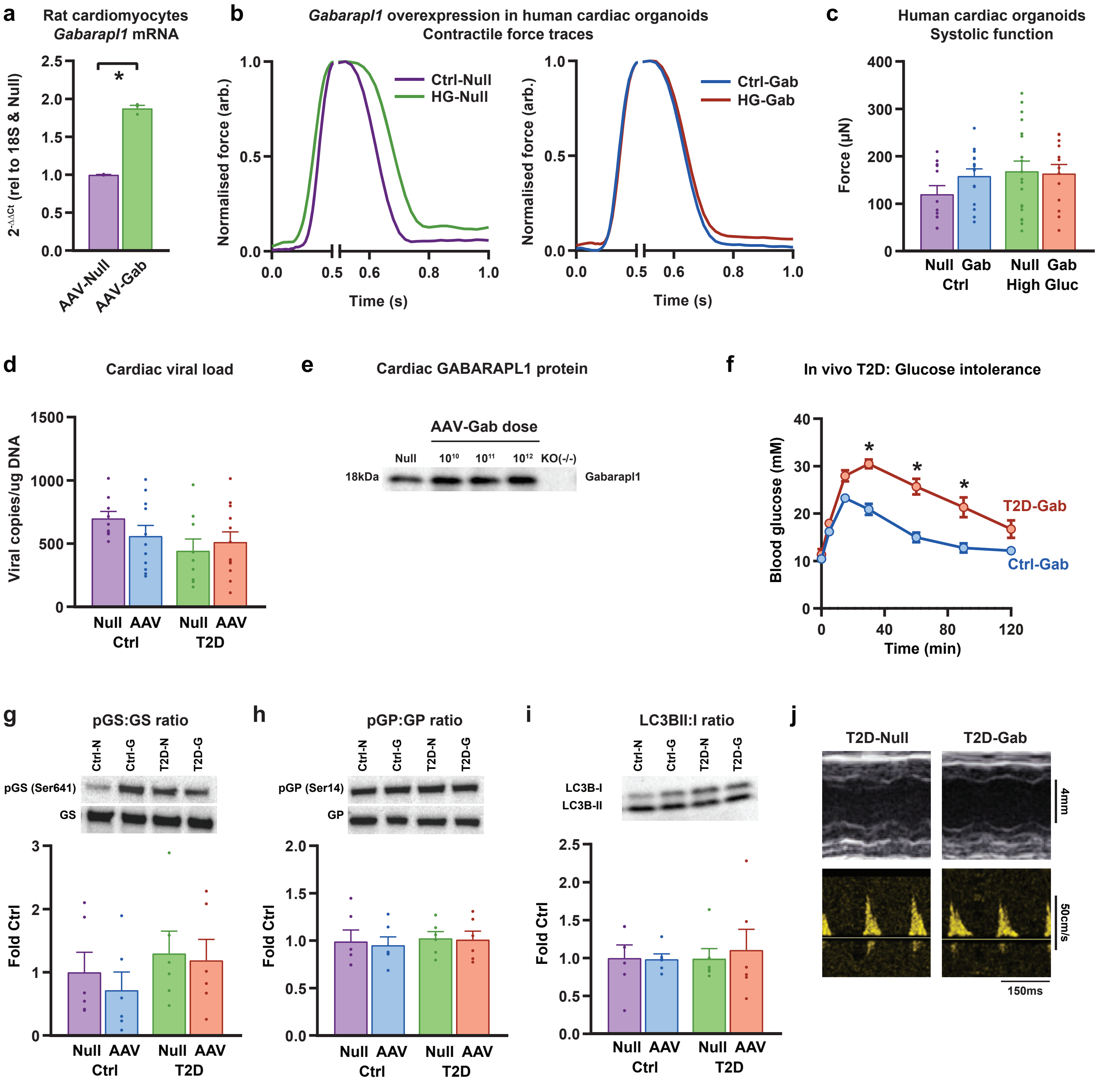


**Figure S5. AAV9-cTnT-*Gabarapl1* in vitro and in vivo cardiac overexpression and characterization of AAV-treated type 2 diabetic (T2D) mice.**
**a,** *Gabarapl1* mRNA expression in neonatal rat ventricular cardiomyocytes transduced with AAV9-cTnT-Null (AAV-Null) or AAV9-cTnT-*Gabarapl1* (AAV-Gab; n=3 independent wells). **b,** Exemplar force traces from human induced pluripotent stem cell-derived cardiac organoids exposed to control glucose (Ctrl, blue) or high glucose (HG, red), transduced with AAV-Null or AAV-Gab. **c,** Force production from human induced pluripotent stem cell-derived cardiac organoids exposed to control glucose (Ctrl) or high glucose (High Gluc), transduced with AAV-Null (Null) or AAV-Gab (Gab), n=12-19 organoids. **d,** AAV vector burden in mouse heart 12 weeks post-injection (i.v.) of $10^{12}$ gc/mouse AAV9-cTnT-Null (AAV-Null) or AAV9-cTnT-*Gabarapl1* (AAV-Gab; digital-droplet PCR detection of the WPRE viral element). **e,** Immunoblot of Gabarapl1 protein expression in mouse hearts 4 weeks post-injection (i.v.) of AAV-Null or AAV-Gab ($10^{10}$, $10^{11}$ or $10^{12}$ gc/mouse) with homozygote *Gabarapl1*-KO mouse heart as negative control. **f,** T2D-induced glucose intolerance is not affected by in vivo cardiac-specific AAV-Gab gene delivery (high fat diet, n=10 mice). **g,** Ratio of phosphorylated to total glycogen synthase (GS) is unchanged in hearts of mice with T2D and *Gabarapl1* overexpression (AAV-Gab, n=6 mice). **h,** Ratio of phosphorylated to total glycogen phosphorylase (GP) is unchanged in hearts of mice with T2D and *Gabarapl1* overexpression (AAV-Gab, n=6 mice). **i,** Ratio of lipidated (II) to unlipidated (I) LC3B protein expression is unchanged in hearts of mice with T2D and *Gabarapl1* overexpression (AAV-Gab, n=6 mice). **j,** M-mode and flow dopper echocardiography exemplar traces in T2D mice treated with AAV-Null or AAV-Gab. Data are presented as mean ± s.e.m.*$p < 0.05$. Related to Figures 5b-c & 5d-i.

**Table S4. AAV9-cTnT-*Gabarapl1* T2D mice: systemic and cardiac characteristics.** T2D mice (high fat diet-fed) injected with AAV9-cTnT-Null (AAV-Null) and AAV9-cTnT-*Gabarapl1* (AAV-Gab) were assessed at 12 weeks post-AAV injection (i.v.) with 28 week diet feeding (n=10 mice). Data are presented as mean ± s.e.m.*p< 0.05. Related to Figures 5d-i.

| | **AAV-Null** | | **AAV-Gab** | |
|---|---|---|---|---|
| | **Ctrl** | **T2D** | **Ctrl** | **T2D** |
| Body weight (g) | 30.4 ± 0.7 | 46.1 ± 1.0* | 31.2 ± 0.7 | 46.5 ±1.4* |
| Blood glucose (mM) | 9.8 ± 0.8 | 12.3 ± 1.1 | 10.0 ± 0.5 | 11.9 ± 0.6 |
| Heart weight (mg) | 145.4 ± 6.0 | 169.1 ± 8.0* | 142.2 ± 3.0 | 171.4 ± 5.6* |
| Heart weight/body weight (mg/g) | 4.78 ± 0.1 | 3.68 ± 0.2* | 4.58 ± 0.1 | 3.70 ± 0.1* |
| Tibia length (mm) | 18.3 ± 0.1 | 18.1 ± 0.1 | 18.2 ± 0.1 | 18.3 ± 0.1 |
| Heart weight/tibia length (mg/mm) | 7.95 ± 0.3 | 9.32 ± 0.4* | 7.82 ± 0.1 | 9.35 ± 0.3* |
| Ejection fraction (%) | 70.7 ± 2.0 | 69.4 ± 1.6 | 66.3 ± 0.9 | 62.8 ± 2.2 |
| Fractional shortening (%) | 31.8 ± 1.1 | 29.8 ± 1.0 | 29.6 ± 0.6 | 26.8 ± 1.0 |
